# FET proteins and PARylation-dependent condensates promote replication fork reversal and genome stability

**DOI:** 10.64898/2025.12.01.691651

**Authors:** Celeste Giansanti, Jack C. Schultz, Jessica Jackson, Alessandro Vindigni, David Cortez

## Abstract

Targeting replication-associated DNA repair mechanisms, including those regulated by PARP1/2 and PARG control of ADP-ribosylation is a powerful cancer therapeutic approach. However, the mechanisms by which PARG inhibition impacts DNA replication remain unclear. Here we combined isolation of proteins on nascent DNA (iPOND) with quantitative proteomics and functional assays to investigate replication fork dynamics upon acute PARG inhibition. We found that FET family proteins (FUS, EWS, and TAF15) are recruited to replication forks in a PAR-dependent manner, forming condensates that slow fork progression and promote fork reversal. FET proteins control fork dynamics in response to some, but not all, replication stresses. FUS inactivation leads to unrestrained fork progression via RECQ1 and PRIMPOL, increased single-stranded DNA gaps, genome instability, and synthetic lethality with BRCA1 deficiency. These findings reveal that FET protein condensates modulate replication stress responses, influencing genome stability and the cellular response to cancer therapeutics targeting PARylation pathways.

## Introduction

Many chemotherapies and radiation treatments for cancer act by inducing DNA damage and disrupting DNA replication, but these approaches often cause significant collateral damage to normal tissues. To address this, considerable effort has been put into studying the mechanisms of DNA repair and replication stress responses that control cancer cell survival with the goal of developing more cancer-specific therapies. One successful strategy is to target DNA damage repair pathways using synthetic lethality, particularly in tumors deficient in homology-directed repair (HR) due to inactivation of genes such as *BRCA1* or *BRCA2* genes^1–3^. In these HR-deficient tumors, agents that interfere with protein poly-ADP-ribosylation (PAR) by PARP enzymes exploit a large therapeutic window, selectively killing cancer cells while sparing normal cells^4,5^.

Protein ADP-ribosylation is a widespread posttranslational modification catalyzed by ADP-ribosyltransferases (ARTs). PARP1, along with PARP2 and PARP3, are involved in DNA replication and damage repair^6–9^. Once bound to DNA breaks, PARP catalyzes PAR chain initiation and elongation^10^. Initiation consists of the transfer of an ADP-ribose moiety from NAD+ to the side chain of an amino acid of a target protein, with the release of nicotinamide; when only a single ADP-ribose is attached, this modification is termed mono(ADP-ribosyl)ation (MARylation). Elongation follows, in which additional ADP-ribose units are polymerized onto the initial ADP-ribose via the ribose 2’-hydroxyl group, resulting in the formation of poly(ADP-ribosyl)ation (PARylation)^11–13^. PAR is rapidly removed by PARG (Poly(ADP-ribose) glycohydrolase), which catalyzes the hydrolysis of the glycosidic bonds between ADP-ribose units of PAR chains^14–16^. PARG has both exoglycosidase and endoglycosidase activities; however, its processivity decreases as PAR chains become shorter, and it cannot remove MARylation^17^. During DNA replication, the processing of Okazaki fragments can activate PARP, leading to the accumulation of PAR chains, which in turn can alter replication fork speed^18,19^. Therefore, PARG activity is essential to remove these PAR chains and maintain normal replication dynamics.

PARP inhibitors are now widely used to treat HR-deficient breast, ovarian, prostate, and pancreatic cancers, although both *de novo* and acquired resistance can limit their effectiveness. Pharmacological inhibitors targeting other DNA repair enzymes, including PARG, are actively being developed as additional cancer therapeutic tools^20,21^.

PARG inhibition preferentially kills cells with DNA strand breaks and those experiencing challenges to DNA replication, such as impaired joining of Okazaki fragments^22–31^. Several PARG inhibitors have been developed, including PDD00017272/3, IDE161, ETX-19477, XNW29016, JA2131, and COH34^23,27,32–35^. Notably, the FDA has granted fast-track designation to IDE161 for platinum-resistant, *BRCA1*/*2*-mutant, advanced or metastatic ovarian cancer, and HR+, HER2-negative breast cancers^36,37^. PARG inhibitors are also being considered for use in HR-deficient breast, endometrial, and gastrointestinal cancers^38^. However, the specific vulnerabilities that determine sensitivity to PARG inhibitors and their precise mechanisms of action remain incompletely understood, and further elucidation of these factors is crucial for optimizing the use of PARG inhibitors in precision oncology.

Multiple pieces of evidence indicate that PARG inhibition exerts its cytotoxicity by interfering with DNA replication. First, PARG inhibitor sensitivity is characterized by persistent DNA replication stress and an accumulation of PAR at sites of replication^22,23,25,39^. Second, the subset of cell lines intrinsically sensitive to PARG inhibition contain defects in genes involved in DNA replication^40^. Furthermore, interfering with many replication factors confers PARG inhibitor sensitivity. Among these are components of the fork protection complex, TIMELESS, TIPIN, and CLASPIN, Okazaki processing factors EXO1 and FEN1, and the major replication stress response kinases ATR and CHK1^22,24,25,39^. Third, PARG travels with the replication machinery, interacts with PCNA, and its localization is regulated by replication stress^41–43^. Fourth, PARG activity prevents the accumulation of abnormal replication structures, and PARG inactivation triggers replication fork stalling and the accumulation of reversed replication forks^23,44^.

To investigate how PAR accumulation interferes with DNA replication and to gain insights into the molecular determinants of PARG inhibitor sensitivity, we used Isolation of Proteins on Nascent DNA (iPOND)^45^ combined with quantitative mass spectrometry^42,46^ to analyze changes in the replication fork proteome following PARG inhibition. iPOND enables the identification of proteins associated with newly synthesized DNA, providing a snapshot of the replication fork environment. As expected, we found that the accumulation of PAR induced by PARG inhibition enhances the recruitment of PAR-binding proteins involved in the base excision repair (BER) and single-strand break repair (SSBR) pathways^39^. Importantly, we also observed that three related proteins, FUS (fused in sarcoma/translocated in sarcoma), EWS (Ewing sarcoma protein), and TAF15 (TATA box-binding protein-associated factor 68 kDa), collectively abbreviated as FET proteins, are among the most enriched proteins on nascent DNA in the presence of the PARG inhibitor.

FET proteins are intrinsically disordered proteins that bind poly(ADP-ribose) (PAR), DNA, and RNA, and are involved in diverse cellular processes including transcription regulation, RNA splicing, transport, and microRNA processing^47–51^. Beyond their roles in RNA metabolism, FET proteins participate in the detection and repair of DNA damage by localizing to sites of laser-induced DNA damage in human cells and facilitating the recruitment of additional DNA damage response factors^52–56^. FUS, a member of the FET family, promotes homologous DNA pairing and is required for efficient double-strand break repair by both homologous recombination and nonhomologous end joining in cancer cells and neurons^54,57,58^. Mutations in FET proteins are associated with pathological protein aggregation in neurodegenerative diseases such as amyotrophic lateral sclerosis (ALS) and frontotemporal lobar degeneration (FTLD)^59–64^, and FET genes are frequently involved in gene translocations that result in fusion proteins in sarcomas^65^.

FET proteins contain two low complexity domains (LCDs): a carboxyl-terminal, positively charged arginine–glycine–glycine (RGG)-rich LCD that acts as a PAR sensor, and an amino-terminal, prion-like SYQG-rich LCD that stabilizes PAR-induced liquid-liquid phase separation, leading to the formation of protein condensates^52,66,67^. In this study, we show that FET proteins and FUS-dependent condensates regulate DNA replication fork progression and fork reversal in response to certain, but not all, replication stress-inducing agents. When FET proteins are inactivated, unrestrained fork progression is mediated by RECQ1 and PRIMPOL. FET proteins are necessary to prevent the accumulation of single-stranded DNA (ssDNA) gaps and to maintain genome stability and cell proliferation, particularly in the context of BRCA1 deficiency.

## Results

### FET proteins are enriched on nascent DNA at replication forks upon PARG inhibition

To characterize how PARG inhibition impacts replisome composition, we utilized iPOND combined with quantitative proteomics using stable isotope labeling by amino acids in cell culture (SILAC) mass spectrometry (MS)^42,45,68,69^ (Fig. 1a). The PAR-binding proteins XRCC1, POLB and LIG3 are among the most enriched proteins on nascent DNA in the presence of the PARG inhibitor PDD00017273 (Fig. 1b; Supplemental Table 1). Those proteins are recruited to sites of DNA damage by PARylation as part of the base excision repair (BER) and single-strand break repair (SSBR) pathways^8^. Their accumulation in the iPOND data is consistent with an accumulation of PAR at sites of DNA synthesis when de-PARylation by PARG is inhibited. Three related proteins, FUS, EWS, and TAF15, collectively abbreviated as FET proteins, were even more enriched (Fig. 1b; Supplemental Table 1). We confirmed that FUS was localized to replication forks upon PARG inhibition using a proximity ligation assay (PLA) (Fig 1c,d). We also confirmed that replication fork localization was completely PAR-dependent as it was abolished by treatment with the PARP1/2 inhibitor Olaparib (Fig. 1e). These results are consistent with and extend previous observations that FUS is recruited to sites of laser-induced DNA damage through an interaction with PAR^53,55,67,70^.

**Figure 1.**
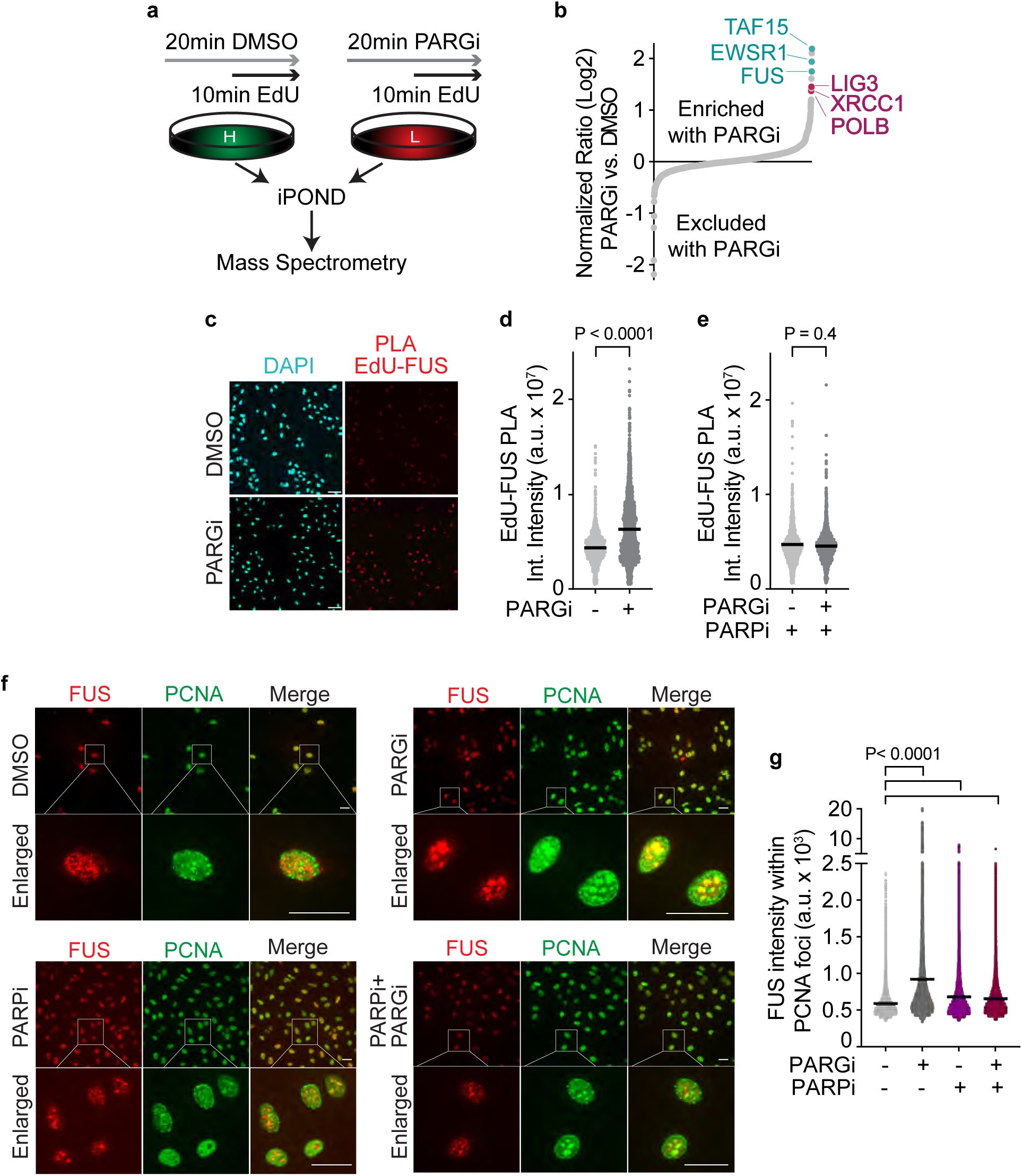
FET proteins are enriched on nascent DNA at replication forks upon acute PARG inhibition. (a) Schematic of iPOND proteomics experiment. HEK293T cells grown with heavy (H) or light (L) amino acids were treated with DMSO or PARG inhibitor (PDD00017273, PARGi, 10 µM) for 20 minutes. EdU was added for the last 10 minutes. Two experiments were completed, swapping the heavy and light cells. (b) Graph of the log2 ratio of PARGi/DMSO for all proteins detected with at least two quantified peptides. (c) Representative images of U2OS cells after PLA assay for EdU and FUS. Cells were treated with DMSO or PARG inhibitor (PDD00017273, PARGi, 10 µM) for 30 minutes. Scale bar, 65 µm. (d) Quantification of PLA assay for EdU and FUS. Individual integrated nuclear intensities per nucleus are plotted. Data are representative of two independent experiments. The black bar indicates the mean. The p-value was calculated using a two-tailed Mann-Whitney U test. (e) PLA assay for EdU and FUS in U2OS cells. In this case, olaparib (10 µM) was added 5 minutes before DMSO or PARGi treatment. (f) Representative images of FUS and PCNA immunofluorescent staining. Scale bar, 30 µm. (g) The average intensity of FUS within PCNA foci was quantified by immunofluorescent staining of U2OS cells after 30 min treatment (PARGi,10 µM; PARPi, 10 µM). A representative experiment from n=2 is shown. One-way ANOVA with Dunn–Šidák’s multiple comparisons post-test was used for statistical analysis.

FET proteins undergo reversible liquid–liquid phase separation to form biomolecular condensates, which coordinate the localization of essential DNA Damage Response (DDR) factors^52^. We analyzed FUS distribution by immunofluorescent staining in U2OS and RPE1-hTERT cells. As expected, nuclear FUS predominantly localizes in discreet foci (Fig. 1f; Supplementary Fig. 1a). Importantly, in PARGi-treated cells, FUS showed increased colocalization with the replication protein PCNA, further corroborating that PAR accumulation triggers FUS foci formation at DNA replication forks (Fig. 1f, g; Supplementary Fig. 1a, b). The intensity of these FUS-PCNA co-localized puncta was greatly reduced by the addition of PARPi (Fig. 1f, g; Supplementary Fig. 1a, b), confirming that the localization depends on PAR. We conclude that FUS and the other FET proteins are recruited to replication forks in cancer (U2OS), transformed (HEK293T), and normal (RPE1-hTERT) cells upon acute PARG inhibition.

### Fork slowing upon PARG inhibition depends on FET proteins and FUS-mediated condensate formation

Previous studies found that PARP activity and persistent PARG inactivation slows replication forks^22,23,25,44,71,72^. The PARG findings were based on RNA interference or long PARG inhibitor treatment times (>24 hours) with a 2.5-hour pre-treatment as the shortest time analyzed^19^. Since PARG inhibition can lead to sequestration of NAD+ and metabolic changes, we wanted to understand the direct effects of acute PARG inactivation on DNA replication. Since FUS inactivation was also previously reported to cause slower replication fork speeds^73^, we also wanted to assess whether FUS and the other FET proteins work in the same pathway with PARG to control replication fork progression. Therefore, we utilized molecular DNA combing with acute PARG inhibitor treatment during the nucleoside analog labeling period (Fig. 2a). Acute treatment of cells with the PARG inhibitor PDD00017273 induced a significant reduction in fork speed, which was dependent on PARP1/2 function since Olaparib abolished this effect (Fig. 2b, c). However, we did not observe altered fork speeds when FUS or the other FET proteins were inactivated by themselves (Fig. 2c). Unexpectedly, inactivating any of the FET proteins individually or all together restored fork speeds to normal in the PARG inhibitor-treated cells (Fig. 2b, c; Supplementary Fig. 2a). The PARG inhibitor treatment increased PAR levels irrespective of FET protein knockdown (Supplementary Fig. 2a, b). Rescue of PARG inhibitor-dependent fork slowing was confirmed in cells where FUS was knocked out by CRISPR-mediated gene editing (Fig. 2d). We also confirmed the same effects using a second PARG inhibitor, COH34^27^ (Fig. 2e). Thus, FET proteins are required to slow replication in response to acute PARG inhibition.

**Figure 2.**
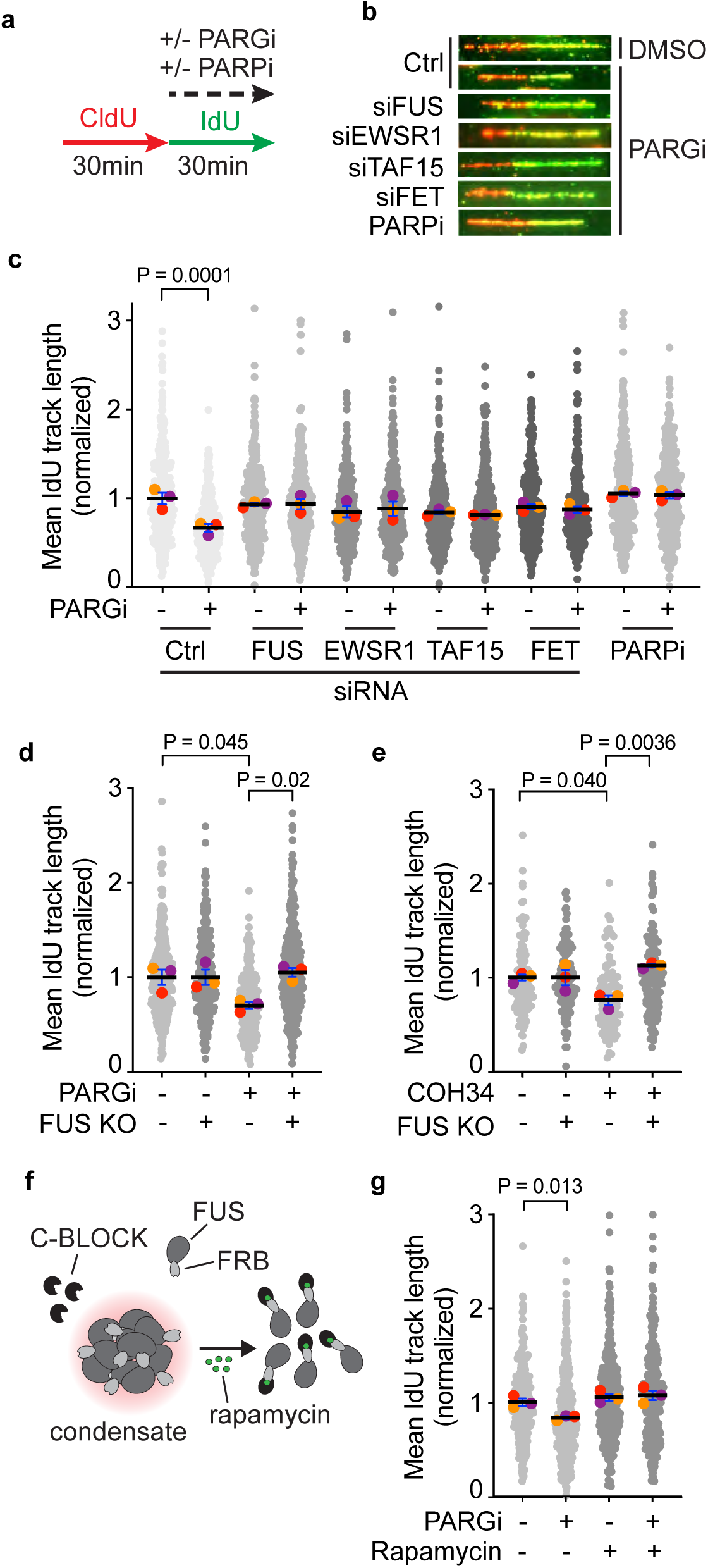
Fork slowing upon PARG inhibition depends on the FET proteins and FUS-mediated condensate formation. Replication fork speed was measured by molecular DNA combing. (a) Treatment and newly synthesized DNA labeling scheme. FET proteins were silenced by siRNAs in HEK293T cells (48 hours), followed by treatment with PARG inhibitor (10 µM) and/or Olaparib (PARPi, 10 µM) as indicated. (b) Representative images of DNA fibers. (c) Quantification of DNA combing results. The mean values from three independent experiments (color circles), along with all individual datapoints (grey) and the mean of the means (black bar), are shown. (d) Synthesized DNA track lengths were measured in control or FUS knockout (FUS KO) HEK293T cells treated with PDD00017273 (PARGi, 10 µM) or (e) COH34 PARG inhibitor (10 µM). (f) Schematic of the DisCo system to disrupt FUS condensate formation^52^. (g) Molecular DNA combing using the DisCo system. 333nM rapamycin was used to cause condensate disassembly during the IdU labeling period, where indicated. In all experiments, values were normalized to the mean value of the untreated control. A minimum of 100 fibers per condition was quantified. P values were derived from analysis of variance (ANOVA) with Dunn–Šidák’s multiple comparisons post-test comparing the mean values. Only comparisons with p-value ≤ 0.05 are displayed.

Next, we examined whether the FET-mediated fork slowing involved the formation of FUS-mediated biomolecular condensates using the DisCo (Disassembly of Condensates) system, which provides inducible control over condensate disruption using rapamycin treatment^74^ (Fig. 2f). Indeed, FUS condensate disassembly completely abrogated fork slowing caused by PARG inhibition, indicating that FUS restrains DNA replication fork progression requires its ability to form biomolecular condensates (Fig. 2g).

### H2O2 and CPT, but not HU, promote FUS enrichment at nascent DNA and fork slowing

We next asked whether the function of FET proteins to slow replication fork elongation was specific to PARG inhibitor-treated cells, or if these proteins are also required to control fork speeds in other replication stress contexts. First, we examined when FUS was enriched at replication forks. We treated cells with hydrogen peroxide (H2O2) to induce oxidative DNA damage, the topoisomerase I inhibitor camptothecin (CPT) that causes topological stress, or hydroxyurea (HU) that both depletes nucleotides and causes oxidative stress by inhibiting ribonucleotide reductase and analyzed FUS localization. In the presence of H2O2, FUS foci appeared smaller than those seen after PARG inhibitor treatment (Fig. 3a compared to Fig. 1f). However, as with PARG inhibitor treatment, these FUS foci often co-localized with PCNA, and the intensity of FUS within PCNA foci was increased in H2O2-treated cells compared to the untreated control (Fig. 3a,b). Similarly, camptothecin strongly enhanced the intensity of FUS within PCNA foci, indicating the re-localization of FUS to DNA replication forks in response to topological stress. In contrast, HU did not affect the FUS signal distribution relative to PCNA (Fig. 3a, b), consistent with a previous iPOND proteomics data that did not observe enrichment of FUS at forks in response to high-dose HU treatment^42^. Notably, unlike H2O2 and CPT, the HU treatment also did not significantly enhance chromatin-bound PAR levels (Supplementary Fig. 2c), suggesting that HU-induced replication stress does not cause as much PARylation as other stresses, and consequently does not induce recruitment of the FET proteins to DNA replication forks.

**Figure 3.**
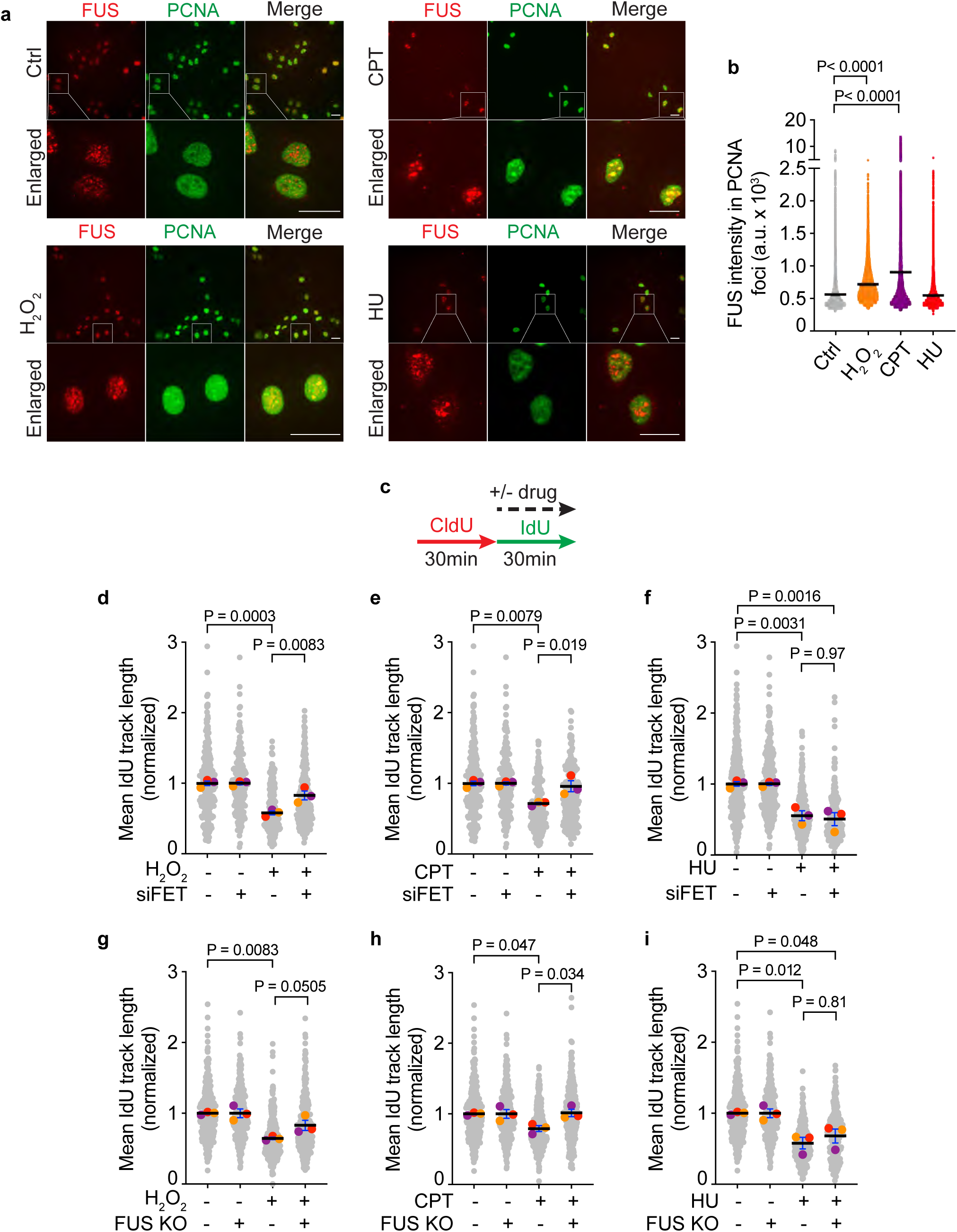
FET proteins are required to slow replication in response to H2O2 and CPT but not HU. (a) Representative images of FUS and PCNA immunofluorescent staining after 30 minutes of treatment with H2O2 (0.1 mM), CPT (100 nM), or HU (100 µM). Scale bar, 30 µm. (b) Quantification of the average intensity of FUS within PCNA foci. One representative experiment out of two is shown. One-way ANOVA with Dunn–Šidák’s multiple comparisons post-test was used for statistical analysis. (c) Replication fork speeds were measured by DNA combing in HEK293T cells in which FUS, EWS, and TAF15 were silenced with siRNAs (d-f) or FUS was knocked out using CRISPR-mediated gene editing (g-i). The mean values from three independent experiments (color circles), along with all individual datapoints (grey) and the mean of the means (black bar), are shown. At least 100 fibers per condition were quantified. One-way ANOVA with Dunn–Šidák’s multiple comparisons post-test comparing the mean values was used for statistical analysis.

To determine whether FET proteins regulate DNA replication fork progression under different types of replication stress, we analyzed DNA synthesis by DNA combing following treatment with H2O2, CPT, or HU (Fig. 3c). As expected, each of these agents induced fork slowing (Fig. 3 d-i). Notably, depletion of FET proteins by siRNA or knockout of FUS prevented fork slowing in response to H2O2 and CPT (Fig. 3d, e, g, h; Supplementary Fig. 2b, d, e), whereas replication fork progression remained reduced in the presence of HU regardless of FET protein status (Fig. 3f, i; Supplementary Fig. 2b, d). Similarly, aphidicolin-induced reduction in fork progression was unaffected by FUS loss (Supplementary Fig. 2e, f). These results indicate that FET proteins are required to slow replication in response to agents such as camptothecin, H2O2, and PARG inhibition, all of which increase PAR levels, but are not required for fork slowing induced by aphidicolin or HU, which directly inhibit DNA polymerases or deplete nucleotide pools.

### FET proteins promote fork reversal to slow DNA replication forks

Replication fork slowing in response to H2O2 and camptothecin is due to fork reversal (Fig. 4a)^74^. Previous studies have shown that genetic inactivation of PARG leads to the accumulation of reversed replication forks^44^. To determine if the FET proteins promote fork slowing in response to acute PARG inhibition by promoting fork reversal, we assessed the requirement for key fork reversal proteins, including RAD51^75^ and the DNA translocases SMARCAL1^76^, ZRANB3^77^, and HLTF^78^, using DNA combing (Fig. 4b). Depletion of RAD51 by siRNA or inactivation of the DNA translocases by CRISPR-mediated gene editing abolished the reduction in DNA synthesis in PARG inhibitor-treated cells (Fig. 4c, d; Supplementary Figure 3a, b), indicating that fork slowing caused by PARG inhibition depends on fork reversal. Notably, co-depletion of FET proteins did not further increase fork progression, suggesting that FET proteins function in the same pathway as fork reversal proteins to mediate fork slowing under replication stress.

**Figure 4.**
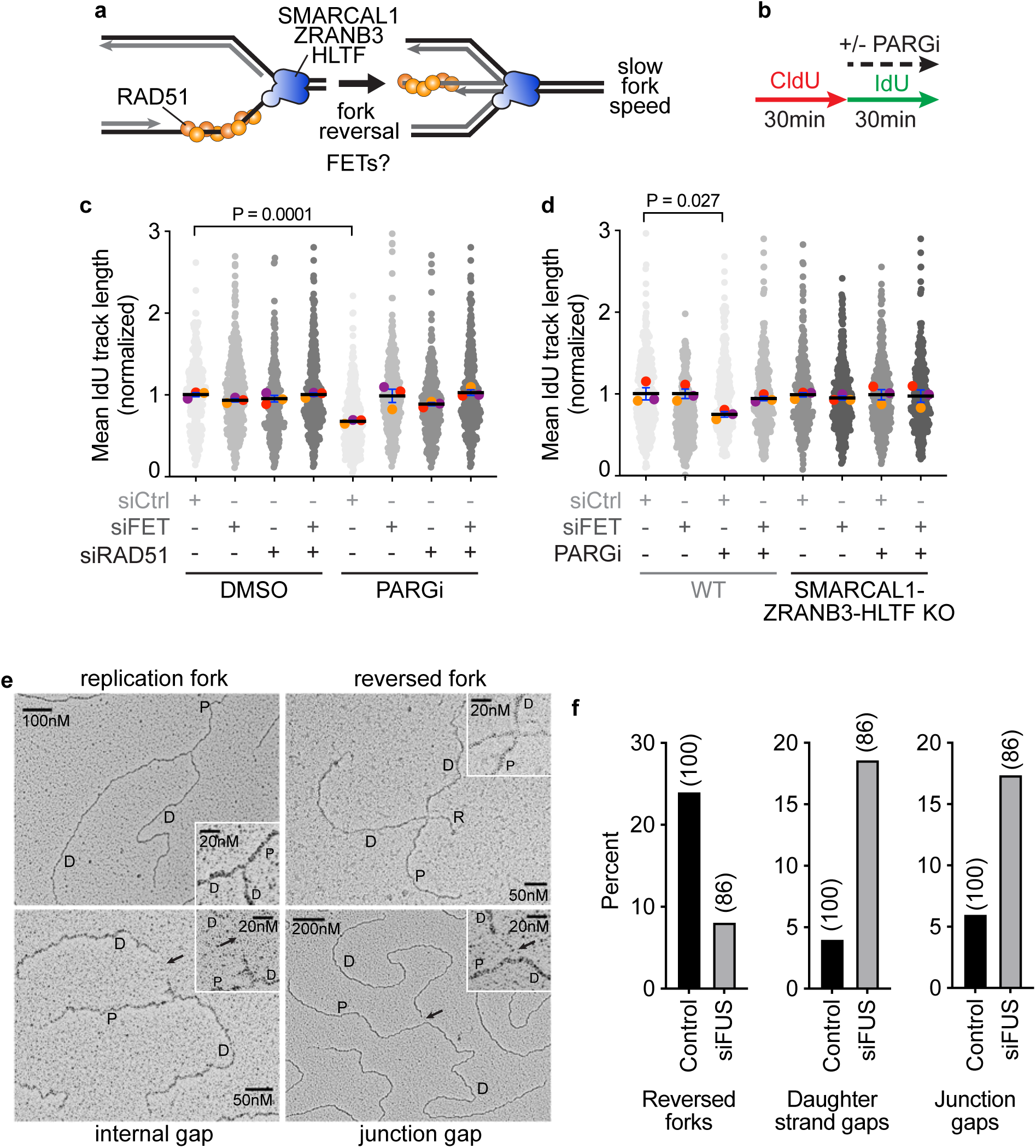
FET proteins promote fork reversal-dependent fork slowing. (a) Schematics of fork remodeling to generate the reversed fork. (b) Replication fork speeds were measured by DNA combing with or without PARGi treatment as indicated. (c) DNA combing in HEK293T cells transfected with siRNAs targeting the FET proteins and/or RAD51. (d) DNA combing using U2OS wild-type or SMARCAL1- ZRANB3-HLTF triple knockout cells with or without silencing of FET proteins with siRNAs. The mean values from three independent experiments (color circles), along with all individual datapoints (grey), and the mean of the means (black bar) are shown. A minimum of 100 fibers per condition was quantified. One-way ANOVA with Dunn–Šidák’s multiple comparisons post-test was used for statistical analysis. Only comparisons with p-value ≤ 0.05 are shown. (e) Examples of replication intermediates caused by 30- minutes of PARGi treatment of U2OS cells. (f) Quantitation of reversed forks and forks with daughter strand or junction gaps after PARG inhibition. 100 replication intermediates were examined from non-targeting siRNA transfected (control) cells. 86 replication intermediates were examined from cells transfected with FUS siRNA.

To confirm that FUS is needed for fork reversal, we examined replication intermediates in wild-type and FUS-deficient cells treated with the PARG inhibitor. As expected, we observed large numbers of reversed forks in the control cells in response to acute PARG inhibition (Fig. 4e,f). These were greatly reduced when FUS was inactivated. Instead, we saw the accumulation of fork structures with ssDNA gaps (Fig 4e,f).

### Unrestrained DNA synthesis in FET-deficient cells depends on RECQ1 and PRIMPOL

The PRIMPOL polymerase can reprime DNA synthesis in response to replication stress, thereby increasing fork speeds but resulting in the formation of ssDNA gaps (Fig. 5a)^79,80^. To determine whether the unrestrained fork speeds and ssDNA gaps observed in FET-deficient cells depend on PRIMPOL-mediated repriming, we used PRIMPOL knockout cells. Loss of PRIMPOL prevented the increase in fork speeds that was otherwise seen upon FET depletion following PARG inhibition (Fig. 5b,c; Supplementary Fig. 4a). Conversely, re-expression of PRIMPOL in these cells restored DNA synthesis rates to normal, indicating that PRIMPOL is responsible for the increased fork speeds when FET proteins are absent. Additionally, inclusion of the ssDNA gap-specific S1 nuclease in the DNA combing assay resulted in a significant reduction in the length of newly synthesized DNA tracks upon FET loss, consistent with the presence of ssDNA gaps due to PRIMPOL activity (Fig. 5d), as also observed in our electron microscopy analyses. These findings indicate that FET proteins promote fork reversal and fork slowing in response to PARG inhibition, thereby preventing PRIMPOL-mediated repriming and the accumulation of ssDNA gaps.

**Figure 5.**
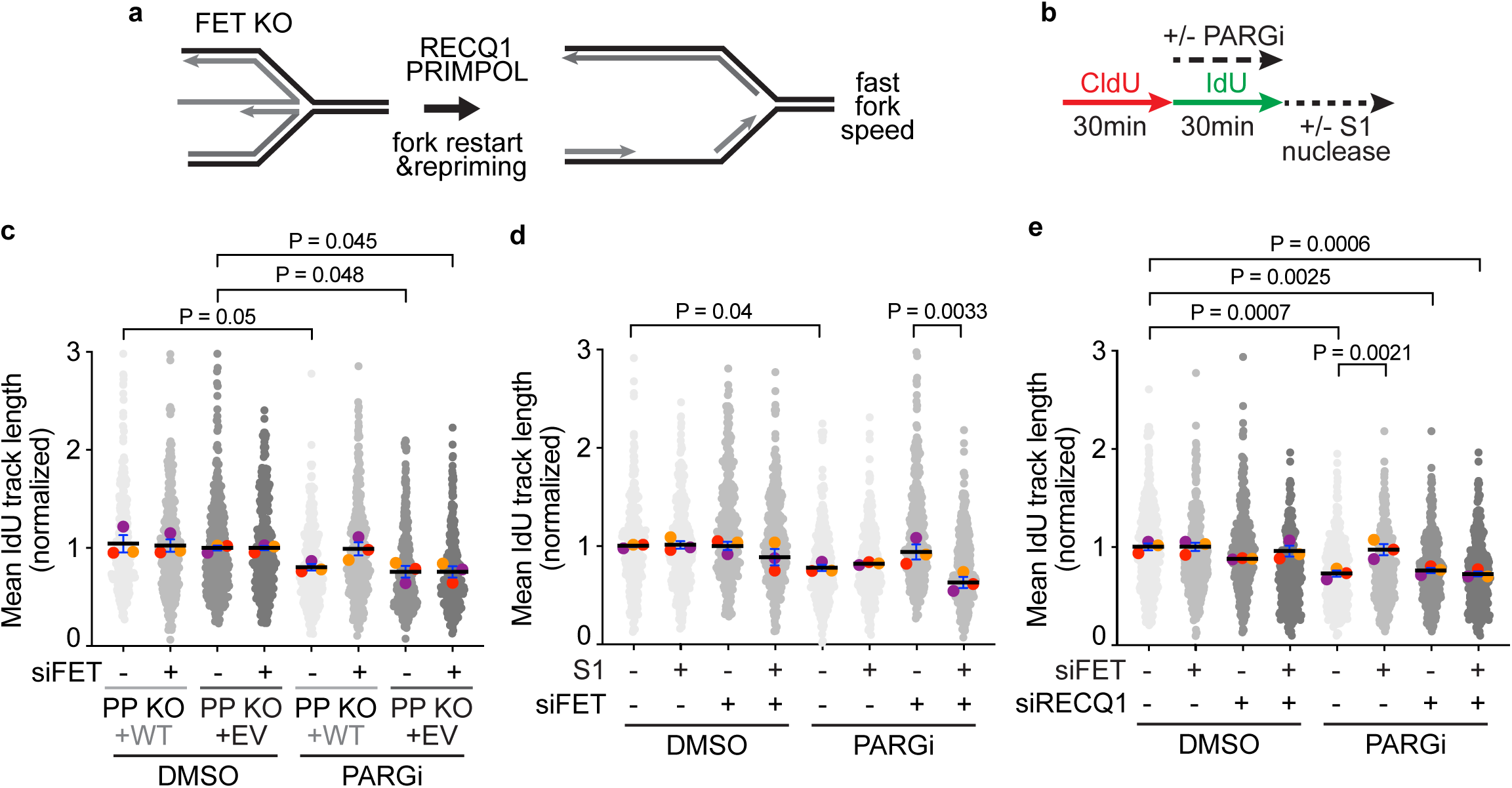
Unrestrained DNA synthesis in FET-deficient cells depends on RECQ1 and PRIMPOL. (a) Schematics of RECQ1- and PRIMPOL-mediated fork restart and repriming, respectively. (b) Newly synthesized DNA track lengths were measured by DNA combing with or without PARGi treatment. (c) DNA combing upon silencing of FET proteins and/or RECQ1 in HEK293T cells. (d) Fork speeds were measured in wild-type or PRIMPOL knockout U2OS cells. (e) Single-stranded DNA gaps created by PARG inhibition when FET proteins are inactivated were measured with DNA combing combined with S1 nuclease digestion, as depicted in b. The mean values from three independent experiments (color circles), along with all individual datapoints (grey), and the mean of the means (black bar) are shown. A minimum of 100 fibers per condition was quantified. One-way ANOVA with Dunn–Šidák’s multiple comparisons post-test was used for statistical analysis.

Previous reports identified a role for the helicase RECQ1 in restarting reversed DNA replication forks and showed that RECQ1-mediated fork restart is regulated by PARylation (Fig. 5a)^81^. Therefore, we used DNA combing to investigate whether the FET proteins antagonized RECQ1 activity, thereby reducing DNA replication fork progression (Fig. 5b). Knockdown of RECQ1 had no effect by itself but restored fork slowing in FET-deficient cells treated with PARG inhibitor (Fig. 5e; Supplementary Fig. 4b), revealing that the unrestrained DNA synthesis observed in FET-deficient cells in the presence of PARG inhibitor depends on RECQ1.

### FUS inactivation causes genome instability and decreased proliferation of BRCA1-deficient cancer cells

The accumulation of gaps on nascent DNA when FET proteins are inactivated prompted us to examine the effects of FUS on genome stability and cellular proliferation. FUS inactivation did not generate a significant hypersensitivity to PARG inhibitors by itself (Supplementary Fig. 5a, b). We also did not observe a significant increase in the DNA damage marker γH2AX in the FUS-deficient cells (Fig. 6a, b; Supplementary Fig. 5c). We reasoned that the detrimental effects of the accumulation of replication-dependent ssDNA gaps on cellular fitness could be limited in HR-proficient cells due to their protection and repair by HR proteins^82,83^. If true, we would expect BRCA1 inactivation to sensitize cells to the loss of FUS and PARG inhibition. Indeed, depleting FUS exacerbated γH2AX accumulation in BRCA1-deficient U2OS cells, and in the breast cancer cell line, HCC1937 (Fig. 6a-c; Supplementary Fig. 5c-e). Importantly, complementation of the BRCA1-defect by re-expression of BRCA1 in HCC1937 prevented the increase in γH2AX (Fig. 6c; Supplementary figure 5e, f). We then analyzed the presence of micronuclei (MN), an indicator of genome instability^84,85^. In control cells, PARG inhibition and FUS inactivation both caused a small increase in micronuclei, and the combined inactivation yielded a small additive effect (Fig. 6d). In BRCA1-deficient cells, inactivating FUS generated a large increase in micronuclei with little or no additional increase when PARG inhibitor was added (Fig. 6d, e).

**Figure 6.**
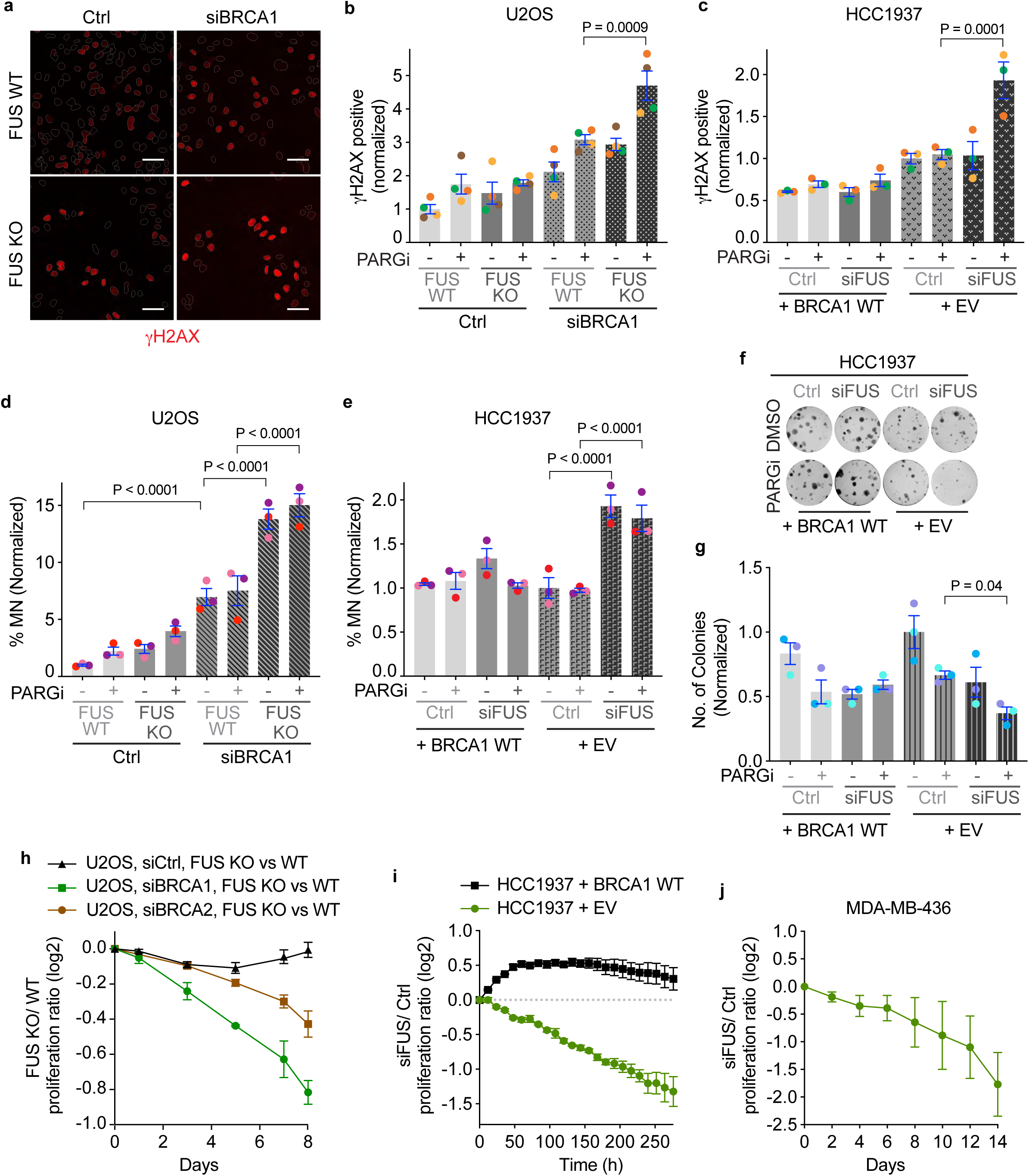
FUS prevents genome instability and maintains cell proliferation in BRCA1-deficient cells. (a) Representative images for γH2AX measured in wild-type (WT) or FUS knock-out (KO) U2OS cells by immunofluorescent imaging 48 h after transfection with the indicated siRNAs, followed by PARG inhibition (10 µM, 24 h). (b) Quantification of the levels of γH2AX in U2OS cells. The mean values from four technical replicates (color circles) are shown. One representative experiment out of five biological replicates is shown. (c) γH2AX quantification in HCC1937 cell line complemented with an empty vector (+ EV) or wild-type BRCA1 (+ BRCA1 WT). The mean values from three technical replicates (color circles) of a representative experiment (n=2) are shown. In b and c, the percentages of γH2AX-positive cells were normalized to the percentage of untreated, wild-type, non-targeting siRNA control. One-way ANOVA with Dunn–Šidák’s multiple comparisons post-test was used for statistical analysis. The percentage of micronuclei (MN)-positive cells in U2OS (d) and HCC1937 (e) was measured by DAPI staining. Three independent experiments are shown as mean ± SEM of replicates. One-way ANOVA with Dunn–Šidák’s multiple comparisons post-test was used for statistical analysis. (f) Representative images of long-term clonogenic assay with parental BRCA1-deficient (right) or the BRCA1-complemented (left) HCC1937 cells. (g) Quantification of colony formation assay. Cells were seeded in low density 24 hours after siRNA transfection and treated with PARG inhibitor (1 μM, 10 days). Three independent experiments are shown as mean ± SEM of replicates. One-way ANOVA with Dunn–Šidák’s multiple comparisons post-test was used for statistical analysis. (h) Competitive cell proliferation assay in PARGi-treated (10uM) wild-type and FUS KO U2OS cells stably expressing either GFP or mCherry. Cells were transfected with a non- targeting control siRNA or an siRNA pool targeting BRCA1 or BRCA2, and data were normalized to the first day of treatment (day 0). Each condition was measured in triplicate wells. Mean ± SEM is displayed. Data are representative of two independent experiments. (i) Competitive cell proliferation assay in PARGi- treated (4 μM) HCC1937 cell line containing an empty vector (HCC1937 + EV) or re-expressing wild-type BRCA1 (HCC1937 + BRCA1 WT). Both cell lines were stably expressing either GFP or mCherry. Data represent the cell count ratio for cells transfected with siRNAs targeting FUS over a non-targeting control siRNA and normalized to the count at time 0. Each condition was measured in quadruplicate wells. Mean ± SEM is displayed. Data are representative of two independent experiments. (j) Competitive cell proliferation assay in PARGi-treated (10uM) MDA-MB-436 cells. Each condition was measured in six technical replicates. Mean ± SEM is shown (n=2).

We then asked how the combination of PARG inhibitor and FUS inactivation would affect cell survival and proliferation of HR-deficient cells. Silencing FUS in the BRCA1-deficient HCC1937 resulted in increased sensitivity toward PARG inhibition in a long-term clonogenic assay (Fig. 6f, g). Again, BRCA1 re-expression restored the cell growth rate to levels comparable to the untreated control (Fig. 6f, g). Similarly, BRCA1 silencing reduced the fitness of FUS knockout compared to wild-type U2OS cells in the presence of PARG inhibitors (Fig. 6h). This effect was not observed in control U2OS cells with an intact HR pathway, and BRCA2 depletion yielded an intermediate phenotype (Fig. 6h). FUS inactivation also restrained cellular proliferation in HCC1937 (Fig. 6i). Complementation of the BRCA1-defect abolished the detrimental effect of FUS loss on cell growth resulting in a small fitness advantage (Fig. 6i). We further confirmed that FUS inactivation reduced cell fitness in two additional BRCA1-mutant breast cancer cell lines, MDA-MB-436, and SUM149PT, and in BRCA1-deficient RPE1-hTERT cells (Fig. 6j; Supplementary Fig. 5h-k). These results indicate that FUS is important for the continued growth and viability of BRCA1- and HR-deficient cellular models.

## Discussion

Replication stress can trigger multiple tolerance pathways, including fork reversal and repriming^86–88^. The balance between these pathways is highly regulated since each can be beneficial or deleterious for genome stability and cell survival, depending on the context^89,90^. Here, we discovered that condensate-forming FET proteins (FUS1, EWS, and TAF15) are important regulators of replication stress response induced by agents that inhibit the PARylation eraser PARG. Specifically, FET proteins shift the balance of response towards replication fork reversal-dependent fork slowing and away from repriming. This activity promotes genome stability by reducing single-stranded DNA gaps. When FET proteins are inactivated or FUS is prevented from forming condensates, the balance shifts towards repriming, leading to increased genome instability and cell death, especially in BRCA1-deficient cell backgrounds. Surprisingly, the FET proteins also regulate these responses when cells are treated with hydrogen peroxide and agents that inhibit topoisomerase I, but not when cells are exposed to polymerase or ribonucleotide reductase inhibitors. Thus, PARG inhibition, H2O2, and CPT, which do not directly interfere with DNA polymerase catalytic activity but do cause the accumulation of PAR at replication forks, slow DNA synthesis in a FET protein- and fork reversal-dependent manner.

PAR accumulation seeds a liquid-liquid phase separation and condensate formation, supported and stabilized by the accumulation of the FET proteins^52,91^. Our data suggest that this condensate formation occurs at stressed replication forks and controls replication dynamics by promoting fork reversal-dependent slowing and inhibiting RECQ1- and PRIMPOL-dependent ssDNA gap formation and unrestrained DNA synthesis. Exactly how the FET-dependent condensate modulates these effects is unclear. PARylation of the TIMELESS protein promotes its degradation and alters fork dynamics^92^, but it is unclear if the FET proteins participate. PAR can inhibit RECQ1 directly, thereby favoring fork reversal^81^. However, this does not seem to be sufficient in cells since we find that PAR accumulates even when the FET proteins are inactivated, but the forks do not slow in these conditions. The FET protein condensate could contribute to the RECQ1 regulation. It could also act as a physical impediment to other replisome activities to favor helicase and polymerase uncoupling to trigger fork reversal^75,93,94^. It is also possible that the condensate recruits or excludes critical replication factors that control the balance of replication stress responses. Finally, although we observe FUS protein localization that is consistent with biomolecular condensate formation and disrupting FUS condensates causes unrestrained synthesis rates, we cannot fully exclude the possibility that some other function of FUS and other FET proteins is important to control replication speeds. Thus, further studies will be needed to fully understand the action of the FET proteins at the stressed replication forks.

Fork reversal is a universal response to replication stress-inducing agents, including H2O2, CPT, aphidicolin, and hydroxyurea^75^. In each of these cases, reversal slows the rate of DNA synthesis. Our data indicated that PARG inhibition also causes slow DNA synthesis and fork reversal, which is dependent on RAD51, SMARCAL1, ZRANB3, and HLTF. Surprisingly, the FET proteins are only required to slow synthesis in response to PARG inhibitor, H2O2, and CPT, but not aphidicolin or hydroxyurea. The underlying difference correlates with the differences in the amount of PAR generated by each of these agents. Another difference is that HU and aphidicolin slow replicative polymerases more directly than the other agents through either polymerase inhibition or depletion of deoxynucleotide precursors. In contrast, the base damage caused by H2O2, the torsional stress and strand breaks caused by CPT, and the elevated PAR levels caused by PARG inhibitors act differently to trigger the uncoupling of replisome activities needed to trigger fork reversal^93,94^. Perhaps the FET-dependent condensates are required to make the uncoupling persistent enough to cause reversal in these cases. In any case, our study indicates that different types of replication stress engage different mechanisms to control fork dynamics.

Our data indicate that FET-mediated fork slowing is required to prevent the accumulation of ssDNA gaps in PARG inhibitor-treated cells due to the repriming activity of PRIMPOL. This was surprising since a previous study found that PARG inhibition caused ssDNA gaps that were independent of PRIMPOL^39^. Differences in methodology and cell type could be important to explain the differences. We examined the effect of an acute 30-minute treatment with PARG inhibitor in contrast to the more prolonged treatment times in the other study which can yield changes in metabolites. In addition, the other study used DNA fiber analyses instead of DNA combing. Observing ssDNA gaps using the fiber method should require gaps on both the leading and lagging template strands. The gaps on the lagging strand could come from a defect in Okazaki fragment processing; however, it is unclear where the gaps on the leading strand template would come from without PRIMPOL repriming. In any case, the gaps we observe in the PARG inhibitor-treated, FET protein-deficient cells do not appear to cause a large problem except when BRCA1 is also inactivated, where they are associated with higher levels of γH2AX and the formation of micronuclei. They also lead to reduced fitness of BRCA1-deficient cells, including breast cancer cell lines consistent with previous studies indicating the sensitivity of these cells to ssDNA gaps^95^.

FUS is frequently altered in cancers. For example, the *FUS* gene on chromosome 3p21.3 is often deleted, the FUS protein is absent in most lung cancers, and its reduction is associated with worse overall patient survival^96,97^. Our findings that FUS is critical for mediating the replication stress response to DNA-damaging chemotherapeutic agents such as topoisomerase poisons and PARG inhibitors, and that it is required for the viability of BRCA1-deficient cancer cells, suggest that FUS status could be an important factor to consider when testing cancer treatment strategies.

While our iPOND-SILAC-mass spectrometry analysis identified FUS, EWS, and TAF15 as amongst the most highly enriched proteins at replication forks when cells are challenged with PARG inhibitors, a previous study using iPOND to study replisome composition in response to PARG inhibition did not identify these proteins^25^. However, that study utilized a very long (24 hours) treatment with PARG inhibitor compared to our acute 20-minute treatment. In addition, they detected 10-fold less proteins by mass spectrometry and failed to detect known PAR binding protein complexes like XRCC1/POLB/LIG3.

In addition to the FET proteins, we found that many additional proteins are enriched or depleted from nascent DNA upon PARG inhibition (Supplemental Table 1). Some, such as those linked to chromatin regulation and DNA end resection, deserve further investigation and could potentially yield insights into genetic vulnerabilities that could be exploited to improve the clinical application of PARG inhibitors. It will also be useful to repeat the replisome proteomics analysis in additional cell types, such as those that are intrinsically hypersensitive to PARG inhibition.

## Methods

### Cell culture

U2OS (ATCC, HTB-96; osteosarcoma, human female origin), HEK293T cells (ATCC, CRL-3216; epithelial, human female origin) and MDA-MB-436 (ATCC, CRM-HTB-26, cells; breast; mammary gland) were grown in Dulbecco’s modified Eagle’s medium (DMEM, 11965092, Thermo Fisher Scientific) supplemented with 10% fetal bovine serum (GeminiBio, S11550). HCC1937 cells (CRL-2336; breast; duct; mammary gland) were grown in ATCC-formulated RPMI-1640 Medium (Thermo Fisher Scientific, A1049101) supplemented with 10% FBS. SUM149PT cells (CVCL_3422, human female triple-negative breast cancer origin) were cultured using Ham’s F-12 Medium (Thermo Fisher Scientific,11765054) supplemented with EGM-2 SingleQuots (Lonza, CC-4176), 10% FBS and 10 mM HEPES (Thermo Fisher Scientific, 15630080). All cell lines were purchased from ATCC, cultured under standard conditions at 37 °C, 5% CO2, and regularly tested for mycoplasma (Universal Mycoplasma Detection kit; ATCC, 30- 1012K).

### RNA interference

U2OS, MDA-MB-436 and HCC1937 cells were transiently transfected with 40 nM siRNA using DharmaFECT 1 transfection reagent (Horizon Discovery Biosciences, 30-1012K), and HEK293T and SUM149PT cells were transiently transfected with 20–25 nM siRNA using Lipofectamine RNAiMax (ThermoFisher Scientific, 13778030) for 48 h, according to the manufacturer’s instructions. siRNAs were obtained from Horizon Discovery Biosciences or Qiagen (FUS, L-009497-00-0005; EWSR1, L-005119-02- 0005; TAF15, L-008930-00-0005; RECQ1, L-013597-00-0005; BRCA1, L-003461-00-0005; BRCA2, SI02653434; RAD51, J-003530-11-0005, J-003530-12-0005; AllStars negative control, 1027281). Target protein depletion was confirmed by immunoblotting.

### iPOND

iPOND, isolation of proteins on nascent DNA, combined with stable isotope labeling of amino acids in cell culture (SILAC) mass spectrometry was performed and analyzed as previously desribed^42,45,68,69^. Briefly, iPOND applies click chemistry to link biotin to the nucleoside analog 5-Ethynyl-2′-deoxyuridine (EdU), which is incorporated into newly synthesized DNA. Light and heavy isotope labeled samples were mixed prior to purification of biotin-coupled DNA and associated proteins using streptavidin-coupled C1 magna beads.

Purified proteins were separated by SDS-PAGE, and gel regions above and below the streptavidin band were excised. Gel regions were washed with 100mM ammonium bicarbonate, and proteins were reduced with 4.5 mM DTT and treated with 10mM iodoacetamide. Gel pieces were destained, and proteins were in-gel digested with trypsin as described previously^42^. After gel extracts were dried, peptides were reconstituted in 0.2% formic acid for analysis by LC-coupled tandem mass spectrometry (LC-MS/MS).

Peptides were loaded onto a C18 reverse phase analytical column using a Dionex Ultimate 3000 nanoLC and autosampler and were gradient-eluted at 350 nL/min using a 100 min gradient. The gradient was as follows: 1-77 min, 2–28 %B; 77-87 min, 28–40 %B; 87–92 min, 40-90 %B; 92-93 min, 90 %B; 93-94 min, 90–2 %B; 94-100 min, 2 %B. Peptides were analyzed using a data-dependent (top 20) method on an Orbitrap Exploris 240 mass spectrometer (Thermo Scientific), equipped with a nanoelectrospray ionization source. The acquisition method consisted of MS1 with an AGC target of 3 × 10^6, followed by 20 MS/MS scan events with an AGC target of 1 x 10^5. The intensity threshold for MS/MS was 1 × 10^4. Dynamic exclusion was enabled with a 20 sec exclusion duration, and HCD collision energy was 30 nce.

For peptide and protein identification, data were analyzed using MaxQuant, version 2.1.4.0^98^. MS/MS spectra were searched against a human database created from the UniprotKB protein database. Precursor mass tolerance was 20 ppm for the first search and 4.5 ppm for the main search. Variable modifications included oxidation of methionine and acetylation of protein N-termini, and carbamidomethylation of cysteines was selected as a fixed modification. Enzyme specificity was set to Trypsin/P, and a maximum of 2 missed cleavages were allowed. The target-decoy false discovery rate (FDR) for peptide and protein identification was set to 1%. A multiplicity of 2 was selected, and Arg10 and Lys8 were selected as heavy labels. The requantify option was enabled. MaxQuant results were filtered to include proteins identified with at least 2 unique peptides. Protein groups identified as reverse hits were removed from the datasets. All reported protein groups were identified with two or more distinct peptides and were quantified with one or more ratio counts.

### DNA combing assay

Cells were plated 24 hours before treatment to reach 50-70% confluence. To label newly synthesized DNA, cells were first treated with 5-chloro-2’-deoxyuridine (CldU, 25 μM, 30 minutes) (Sigma-Aldrich, C6891) and then 5-iodo-2’-deoxyuridine (IdU, 25 μM, 30 minutes) (Sigma-Aldrich, l7125). As indicated, treatments with PARG inhibitor (PDD 00017273, Biotechne, 5952) and PARP inhibitor (Olaparib, Selleckchem, S1060), H2O2 (hydrogen peroxide, Millipore Sigma, H1009), HU (hydroxyurea, Millipore Sigma, H8627), CPT (camptothecin, Millipore Sigma, C9911) were added. Approximately 400,000 cells were embedded in 1.5% low-melting agarose plugs in phosphate-buffered saline (PBS) and digested overnight in 0.1% sarkosyl, proteinase K (2 mg/ml), and 50 mM EDTA (pH 8.0) at 50°C. Plugs were washed in TE (10mM Tris pH=8.0, 1mM EDTA), transferred to 100 mM MES (pH 5.7), melted at 68°C, and digested with 1.5 U of β-Agarase I (New England Biolabs, M0392S) overnight at 42°C. DNA was combed on silanized coverslips using a GenomicVision combing apparatus. The DNA was stained with antibodies recognizing IdU (BD, 347580) and CldU (Abcam, ab6326) for 1 hour, washed in PBS, and probed with fluorescent secondary antibodies (Alexa Fluor 488, A28175, Alexa Fluor 555, A-21434) for 45 min. Images were obtained using the Axio Scan.Z1 (Zeiss). Images were manually analyzed for fiber length using Fiji.

### DNA molecular combing with S1 nuclease assay

The DNA molecular combing assay was performed as described above. S1 nuclease (10 U, 2 hours at 37°C) was added to the agarose plugs after TE washes and transfer into 1× S1 nuclease buffer (Thermo Fisher Scientific, EN0321), in the presence of protease inhibitor cocktail (Roche, 05892970001) to inhibit residual proteinase K. Plugs were washed in TE buffer once and then transferred to 100 mM MES (pH 5.7) before continuing the DNA molecular combing protocol.

### Immunofluorescence

Cells were plated in a 96-well clear-bottom plate (Cellvis, P96-0-N) and treated as described, before pre-extraction with 0.4% Triton X-100 in cold PBS on ice for four minutes, followed by fixation with 4.0% formaldehyde solution (prepared from Millipore Sigma, F1635) for 15 minutes at room temperature, and blocking for 1 h at room temperature (10% goat serum, 0.1% Triton X-100 in 1X PBS). Primary antibody incubations with γH2AX (1:1000; Millipore Sigma, 05-636) were performed at 4 °C overnight in blocking buffer. After incubation with fluorescently labeled secondary antibody (Alexa Fluor 488, A28175), nuclei were stained with DAPI (Sigma, D9542). Plates were imaged on a Molecular Devices ImageXpress Micro, and the nuclear integrated intensity was quantified using the MetaXpress Multi Wavelength Cell Scoring software application module.

### Proximity Ligation Assay (PLA)

PLA was performed according to the manufacturer’s instructions using the Duolink PLA kit (Millipore Sigma, DUO92008), *in situ* PLA probe anti-mouse MINUS (Millipore Sigma, DUO92004), and *in situ* PLA probe anti-rabbit PLUS (Millipore Sigma, DUO92002). The protocol was used with the following additions. Cells were plated, pre-extracted and fixed as stated in the immunofluorescence method. Cells were labeled with 10 μM EdU for 30 minutes and at the same time with the drug of interest, when indicated. Then the cells were blocked with 3% BSA and processed via click chemistry with biotin-azide. Biotin antibody (Cell Signaling Technology, 5597) (1:200 dilution) and FUS antibody (Bio-Techne, NBP2-52874) (1:300 dilution) were used. Nuclei were stained with DAPI (Sigma, D9542). Plates were imaged on a Molecular Devices ImageXpress Micro, and the nuclear integrated intensity was quantified using the MetaXpress Multi Wavelength Cell Scoring software application module.

### Immunoblotting

Whole-cell lysates were generated using lysis buffer containing 50 mM Tris (pH 7.4), 200 mM NaCl, 1% IGEPAL CA-630 (Millipore Sigma, I8896), 1 mM EDTA (pH 8.5), Protease Inhibitor Cocktail (Roche, 05892970001), 25 U of Pierce Universal Nuclease (Thermo Fisher Scientific, 88700), 1 mM dithiothreitol (DTT) and 1 mM MgCl2. Whole cell lysates were collected by scraping and incubating cells on ice for 10 minutes with lysis buffer. Equal protein weights were calculated using the DC protein assay kit I (Bio-Rad, 5000111), separated using SDS-PAGE gel and immunoblotted on Amersham Hybond PVDF membrane (10600023). The membrane was blocked in 5% (w/v) non-fat milk in TBS containing 0.1% Tween-20 (Millipore Sigma, P1379) for 1 h and incubated with the primary antibodies at 4°C overnight. For protein detection, the primary antibodies targeting the following proteins were used: FUS (Bio-Techne, NBP2- 52874); EWSR1 (Thermo Fisher Scientific, MA5-24791); TAF15 (Thermo Fisher Scientific, MA5-47431); PRIMPOL (J. Mendez^80^); RAD51 (Abcam, ab213); GAPDH (Proteintech, 60004-1-Ig); γH2AX (EMD Millipore, 05-636); ZRANB3 (Proteintech, 23111-1-AP); HLTF (Abcam, ab183042); SMARCAL1 (Bethyl Laboratories, A301-616A); BRCA1 (Thermo Fisher Scientific, MA1-23164); BRCA2 (Thermo Fisher Scientific, MA5-60184); Poly/Mono-ADP Ribose (Cell Signaling Technology, 89190S). Membranes were then incubated with peroxidase (HRP)-conjugated secondary antibodies (Thermo Fisher Scientific, anti-mouse, 31457, anti-rabbit, 31460) or StarBright Blue 700 Goat Anti-Mouse IgG (Bio-Rad, 12004158). The signal from the HRP-conjugated secondary antibodies was detected by SuperSignal West Pico PLUS Chemiluminescent Substrate (Thermo Fisher Scientific, 34580).

### Two-Color Competition Assay

Cells were transduced with retroviral vectors to generate stable cell lines expressing mCherry or GFP. These cells were mixed in a 1:1 ratio to seed on 96-well clear-bottom plate (Cellvis, P96-0-N) at a density of 3,000 cells per well for experiments. 48 hours after transfection (Day or Time 0), cells were treated with PARG inhibitor or left untreated. The medium with and without PARG inhibitor was refreshed every 2 days. Imaging, analyses for segmentation, and counting of GFP and mCherry positive cells were performed using Molecular Devices ImageXpress Micro and the MetaXpress Multi Wavelength Cell Scoring software application module. Alternatively, imaging and analysis was completed with an Agilent Cytation 5 instrument and Gen5 software. Each experiment was conducted in at least two biological replicates (independent siRNA transfections), and each condition was assessed in at least three technical replicates.

### Viability assays

Viability assays were completed with alamarBlue HS Cell Viability Reagents (Thero Fisher Scientific, DAL1025). Readings were taken at 590 nm 1 h after adding the reagent to the cells. For long term clonogenic survival cells were seeded at low density and allowed to grow for two weeks prior to scoring with methylene blue staining (48% methanol, 2% methylene blue, 50% water). All clonogenic survival assays were completed in triplicate.

### Electron microscopy

Electron microscopy experiments were performed as previously described^99^. DNA was cross-linked by incubating with 10 μg/mL 4,5′,8-trimethylpsoralen (MedChemExpress, HY-B1157), followed by a 3-minute exposure to 366 nm UV light on a precooled metal block, for a total of three rounds. Cells were lysed, genomic DNA was isolated from the nuclei by proteinase K digestion, chloroform-isoamyl alcohol extraction, followed by genomic DNA purification by isopropanol precipitation and digested with PvuII HF with the appropriate buffer for 4 hours at 37°C. Samples were spread on a carbon-coated grid in the presence of benzyl-dimethyl-alkylammonium chloride, followed by platinum rotary shadowing. For imaging, a JEOL JEM-1400 electron microscope using a bottom-mounted AMT XR401 camera was used. Analysis was performed using ImageJ software (National Institute of Health). DNA secondary structures were scored according to the following analysis settings: fibers with a minimum thickness of 10 nm were counted as duplex DNA, fibers with reduced thickness (5-7 nm), as single-stranded DNA (ssDNA). The joining of three DNA fibers into a single junction, with two symmetrical daughter strands and a single parental strand, was counted as a canonical replication fork (three-way junction). Structures with four DNA fibers joined at a single junction, consisting of two symmetrical daughter strands, one parental strand and the addition of a typically shorter fourth strand, representative of the reversed arm, were counted as reversed replication forks (four-way junction). The length of the two daughter strands corresponding to the newly replicated duplex must be equal (b = c), whereas the length of the parental and regressed arms can vary (a ≠ b = c ≠ d). This allows us to distinguish reversed replication forks from canonical Holliday junction structures, as the latter will be characterized by arms of equal length (a = b, c = d). Moreover, the junction of the reversed replication fork must present a bubble structure, indicating that the junction is open and isn’t the result of the occasional crossover of two DNA molecules.

### Statistics and reproducibility

Statistical analysis for all data was completed using Prism 10 (GraphPad). The details of the individual tests are indicated in the respective figure legends. Statistical methods were not used to estimate sample size or to exclude/include datapoints or samples. Multiple approaches were used to verify the results of every experiment to ensure that off-target effects or clonal variations did not cause the results. All experiments were performed at least three times unless otherwise noted in the figure legend. Graphs display all data points from all replicate experiments and the mean of each replicate experiment. Statistical analyses were done using the means of the experiments, not individual data points unless indicated otherwise in the figure legend.

## Data Availability

The proteomics data is available via PRIDE accession number PXD069827. The uncropped immunoblots are available in the supplemental figures.

## Supporting information

Uncropped western blots

## Acknowledgements

We thank Dr. Juan Mendez for the PRIMPOL antibody and Dr. Chandra Tucker for reagents to use the DisCo system. This work was supported by grants to D.C. from the Breast Cancer Research Foundation (BCRF-22-029) and the National Cancer Institute (R01CA239161). Additional support came from the Vanderbilt-Ingram Cancer Center Support Grant (P30CACA068485) and pilot funding from the Dunham Family Fund. C.G. is supported by an American Cancer Society Postdoctoral Fellowship (PF-24-1252799-01). Proteomics was completed in the Vanderbilt Mass Spectrometry Research Center Proteomics Core Laboratory.

## Author Contributions

D.C. and C.G. conceived of the project. D.C. supervised the project. C.G., J.S. performed most experimentation. J.J. performed the electron microscopy with supervision from A.V. C.G. and D.C. wrote the manuscript.

