## Supplemental Figures for "FET proteins and PARylation-dependent condensates promote replication fork reversal and genome stability"

**Contents**

Supplemental Figures 1-5

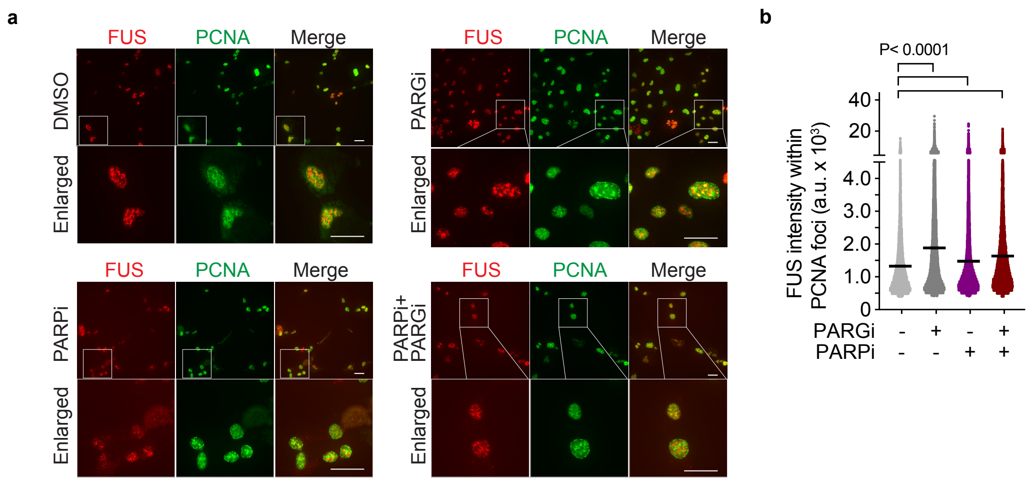


**Supplementary Figure 1. FET proteins are enriched at replication forks upon PARG inhibition.** (a) Representative images of FUS and PCNA immunofluorescent staining in RPE cells after 30 min treatment (PARGi,10 μM; PARPi, 10 μM). Scale bar, 30 μm. (b) The average intensity of FUS within PCNA foci was quantified. One representative experiment out of two is shown. Black bars indicate the means. One-way ANOVA with Dunn–Šidák's multiple comparisons post-test was used for statistical analysis.

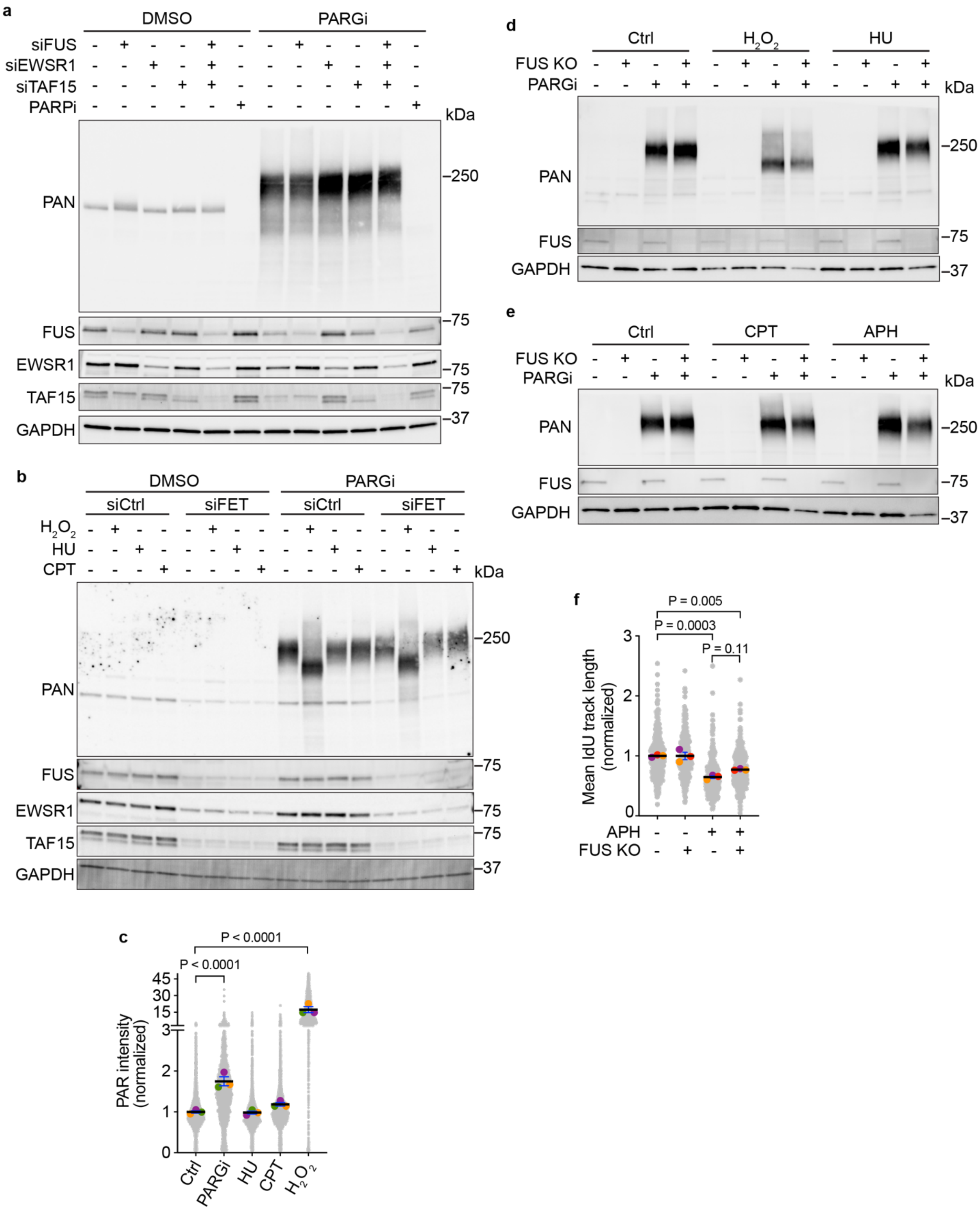


**Supplementary Figure 2. FET proteins regulate DNA replication fork progression.** (a) Immunoblot corresponding to the DNA combing experiment presented in Fig. 2c. FUS, EWSR1, TAF15, and

Poly/mono-ADP-ribose (PAN) are stained to reveal the siRNA transfection efficiency and ADP-ribosylation accumulation in the presence of PARG inhibitor. GAPDH was used as a loading control. (b) Immunoblot corresponding to the DNA combing experiment shown in Fig. 3 d-f. FUS, EWSR1, and TAF15 were stained to test the siRNA transfection efficiency. PAN signal was also measured. PARG inhibitor (10 μM, 30min) combined with the indicated DNA-damaging agents revealed ADP-ribosylation levels across the different treatments. (c) Immunofluorescence was used to quantify the chromatin-associated levels of Poly/mono-ADP-ribose. U2OS cells were treated for 30 minutes with PARGi (10 μM), $H_2O_2$ (0.1 mM), CPT (100 nM), or HU (100 μM). The mean values from three technical replicates (color circles) and the single datapoints (grey circles) are shown. One representative experiment out of two is shown. One-way ANOVA with Dunn–Šidák's multiple comparisons post-test was used for statistical analysis. (d, e) Immunoblot corresponding to the DNA combing experiment shown in Fig. 2d, 3g-i, Supplementary Figure 1f. (f) Synthesized DNA track lengths were measured in control or FUS knockout (FUS KO) HEK293T cells treated with aphidicolin (APH, 2 μM, during IdU labeling). The mean values from three independent experiments (color circles), along with all individual datapoints (grey), and the mean of the means (black bar) are shown.

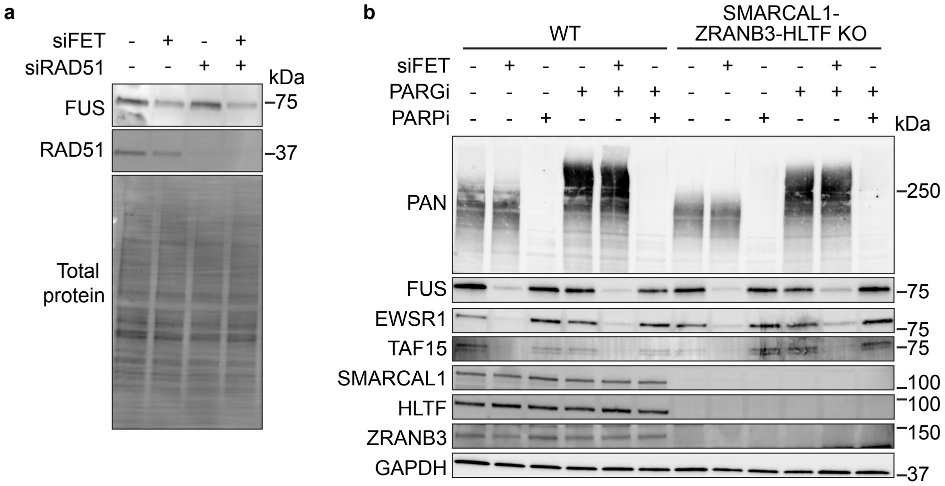


**Supplementary Figure 3. Fork slowing in response to PARG inhibition is dependent on RAD51 and fork reversal enzymes.** (a) Immunoblot corresponding to the DNA combing experiment presented in Figure 4c. Bio-Rad Stain-Free total protein was used as a loading control. (b) Immunoblot corresponding to the DNA combing experiment shown in Fig. 4d.

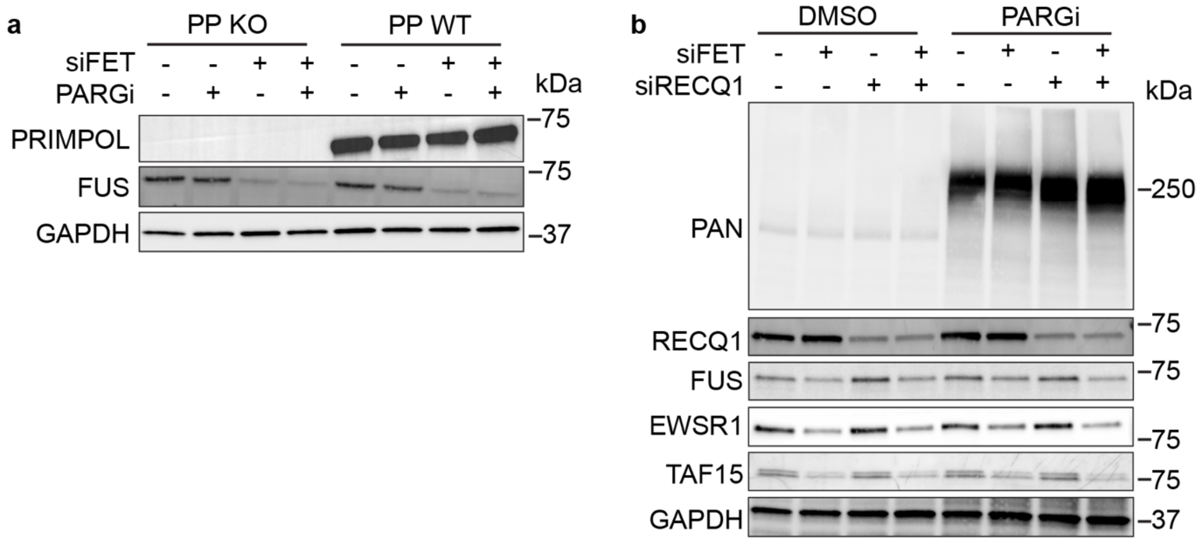


**Supplementary Figure 4. Unrestrained DNA synthesis in FET-deficient cells depends on RECQ1 and PRIMPOL.** (a) Immunoblot corresponding to the DNA combing experiment shown in Fig. 5c. (b) Immunoblot corresponding to the DNA combing experiment presented in Fig. 5e.

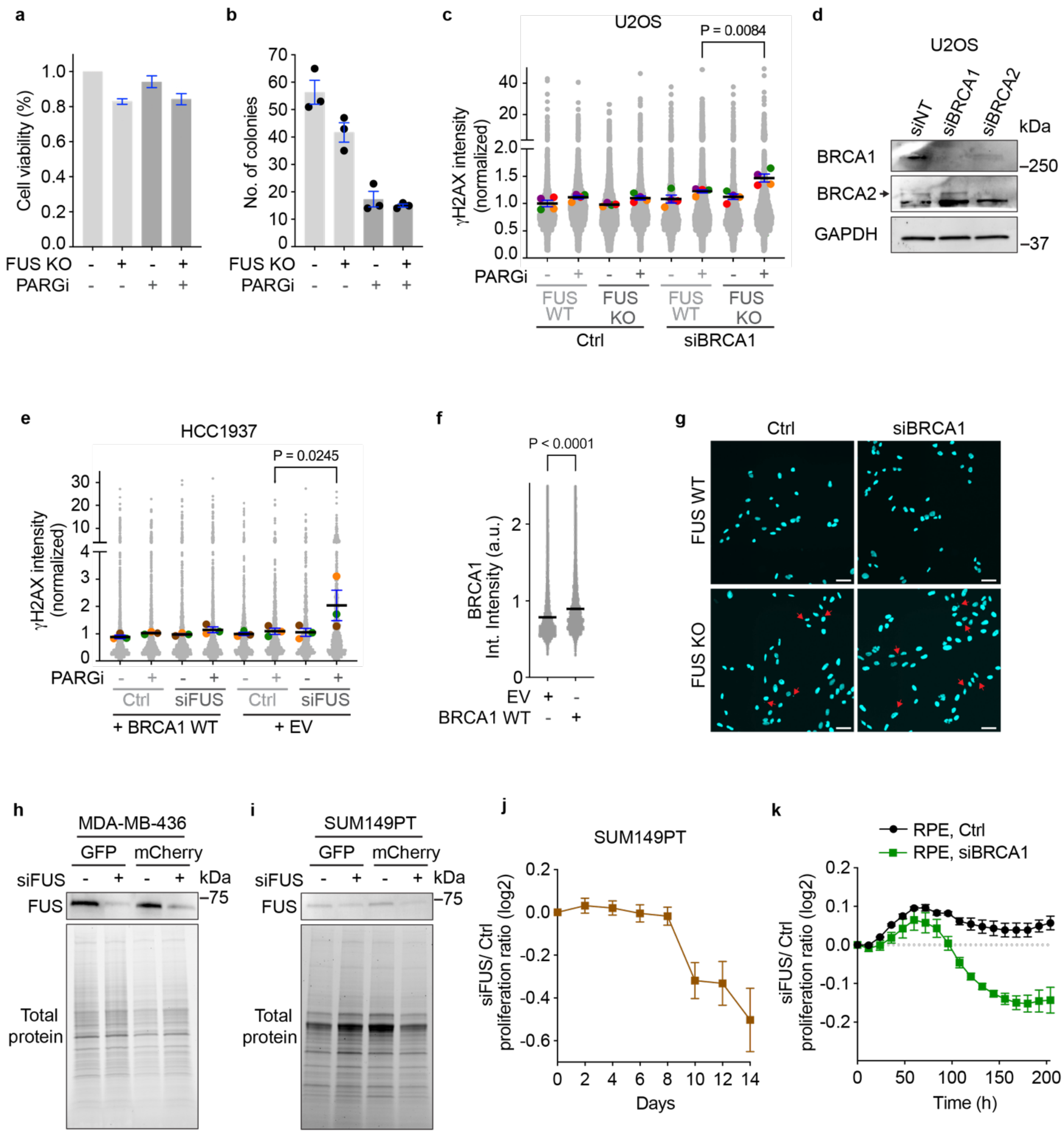


**Supplementary Figure 5. FUS promotes cellular proliferation in BRCA-deficient cells.**

(a) Cell viability assay completed with alamarBlue (Thermo Fisher Scientific). Wildtype (WT) or FUS knockout (KO) U2OS cells were treated with PARGi (10 μM, 8 days). All viability measurements are presented as a percentage of the untreated control. Each bar represents the mean of nine technical replicates. Data are representative of two independent experiments. (b) Colony formation assay. U2OS cells were plated for long-term clonogenic survival. Colonies were allowed to grow for two weeks prior to scoring with methylene blue staining. Bars represent the mean ± SEM values from three independent experiments (black circles). (c) Quantification of γH2AX intensities measured by immunofluorescent staining corresponding to Fig. 6b. Values were normalized to the mean value of the untreated, non-targeting, WT control. The mean values from four technical replicates (color circles), along with all individual datapoints (grey), and the mean of the means (black bar) are shown. One representative experiment out of five is shown. P values were derived from analysis of variance (ANOVA) with Dunn–Šidák's multiple comparisons post-test comparing the mean values. Only comparisons with p-value ≤ 0.05 are displayed. (d) Immunoblot corresponding to competitive proliferation assay in Fig. 6h. BRCA1 and BRCA2 were stained to test siRNA-mediated transfection efficiency in U2OS cells 48 h after transfection. (e) Quantification of γH2AX intensities measured by immunofluorescent staining corresponding to Fig. 6c. (f) Immunofluorescent staining of BRCA1. HCC1937 cells were transduced with an empty vector (EV) or an expression vector for the re-expression of wild-type BRCA1. (g) Representative images of DAPI-stained nuclei of wild-type (WT) or FUS knock-out (KO) U2OS cells 72 h after transfection with siRNA pools targeting the indicated factors and PARG inhibitor treatment (10 µM, 24 h). Red arrows indicate micronuclei (MN). (h, i) Immunoblot corresponding to the proliferation assay presented in Fig. 6j, Supplementary Fig. 5j. FUS was stained in MDA-MB436 (h) and SUM149PT (i) to test transfection efficiency. Bio-Rad Stain-Free total protein was used as a loading control. (j) Competitive cell proliferation assay in PARGi-treated (10 µM) SUM149PT cells. Each condition was measured in seven technical replicates. Mean ± SEM is shown (n=2). (k) Competitive cell proliferation assay in PARGi-treated (1 µM) RPE cell line. Data represent the cell count ratio for cells transfected with siRNAs targeting FUS over a non-targeting control siRNA and normalized to the count at time 0. Each condition was measured in quadruplicate wells. Mean ± SEM is displayed. Data are representative of two independent experiments.
