## Supplementary material for "FET proteins and PARylation-dependent condensates promote replication fork reversal and genome stability": Uncropped western blots

### Supplementary Figure 2d

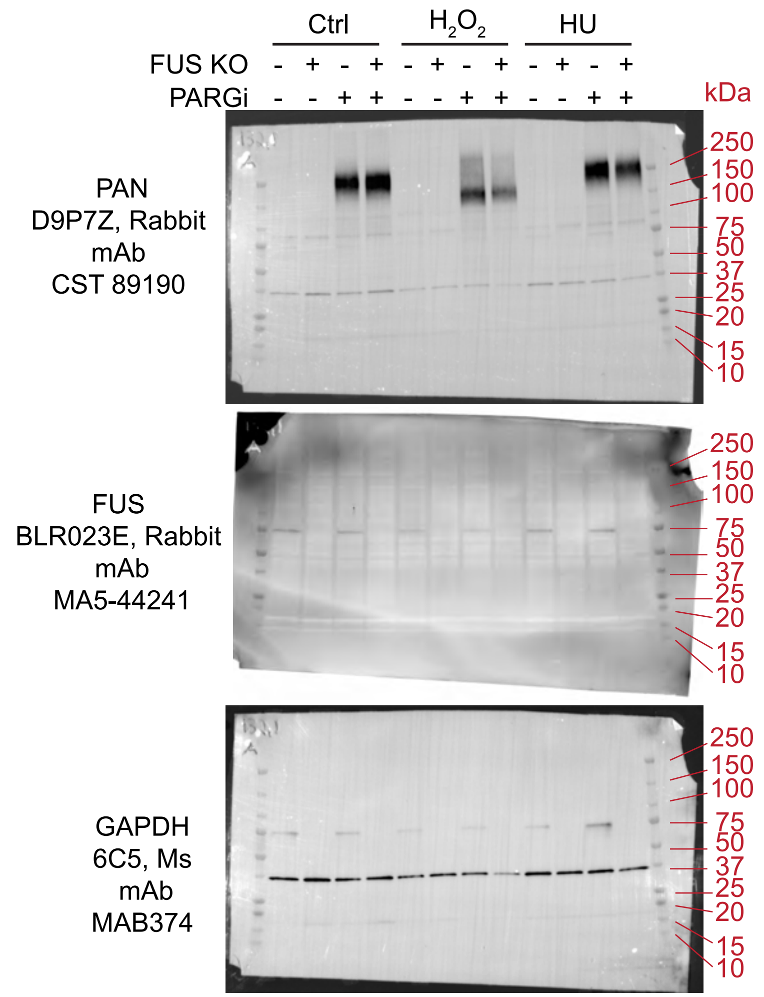


### Supplementary Figure 3a

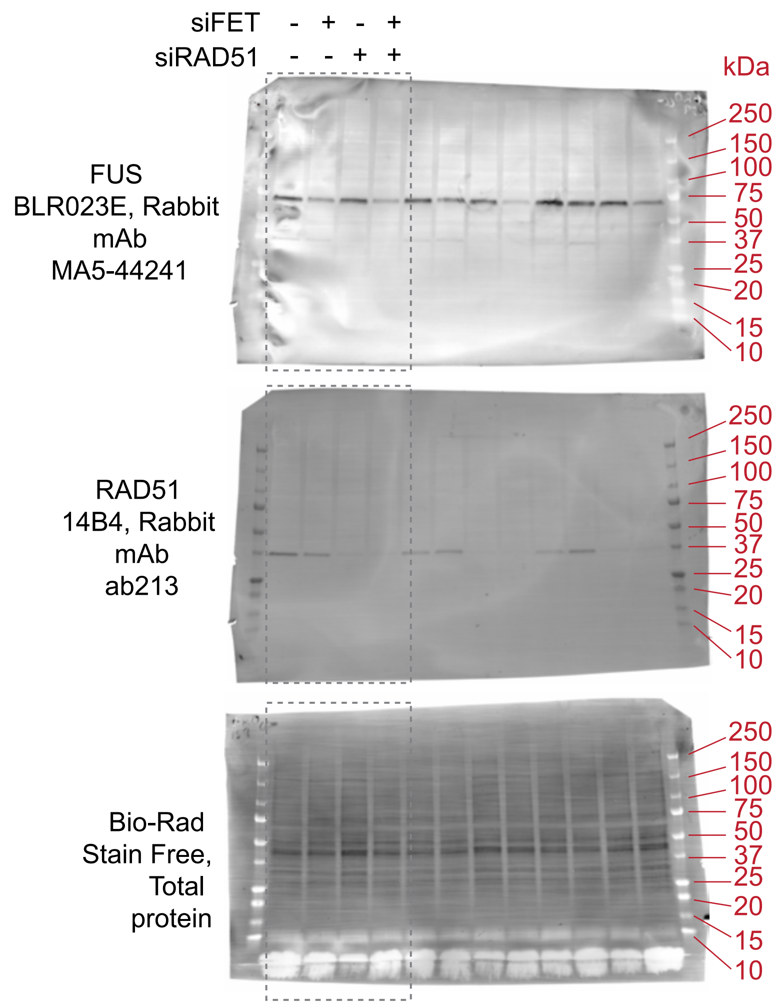


### Supplementary Figure 2e

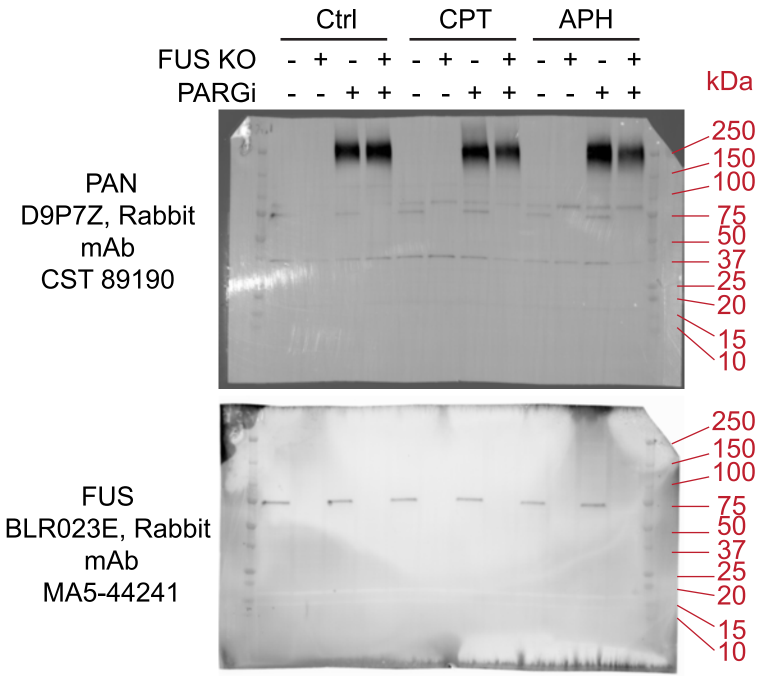


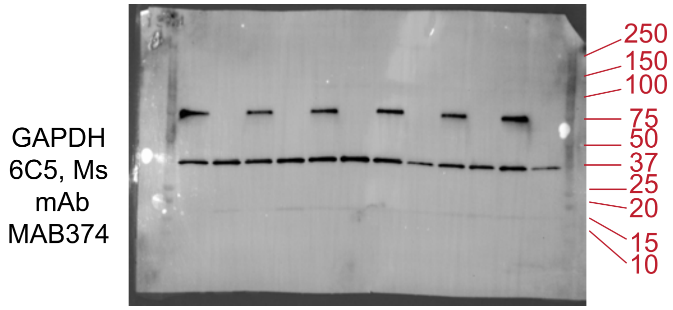

Supplementary Figure 3b

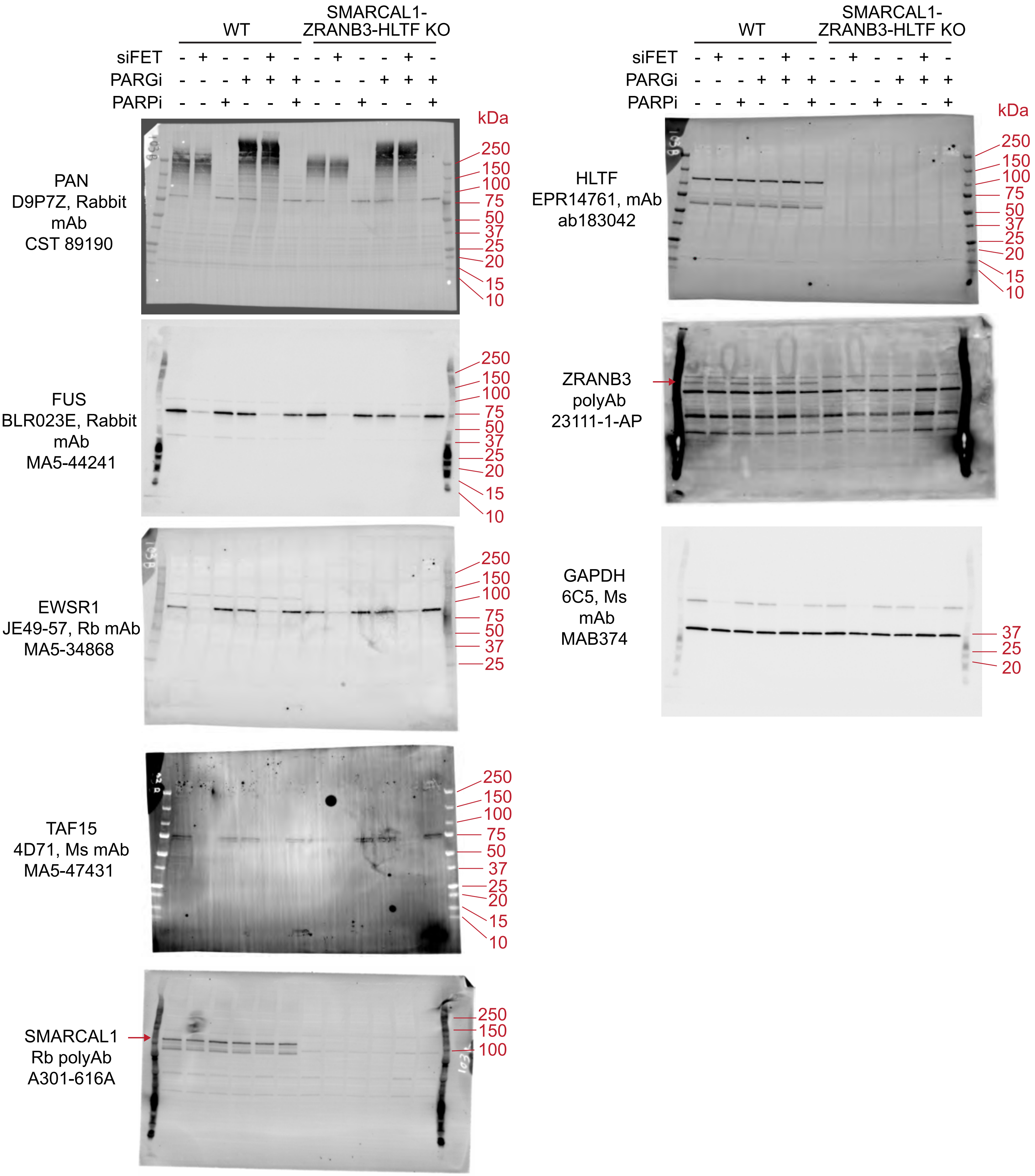

Supplementary Figure 4a

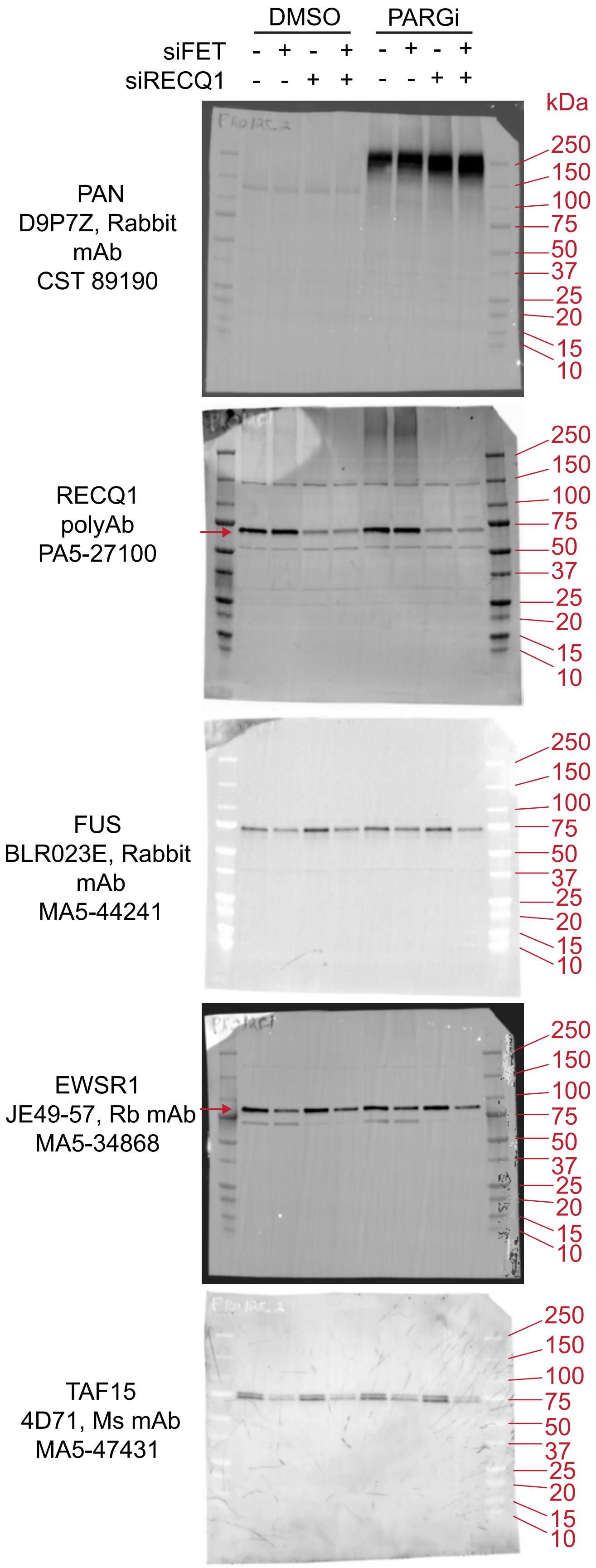


Supplementary Figure 4b

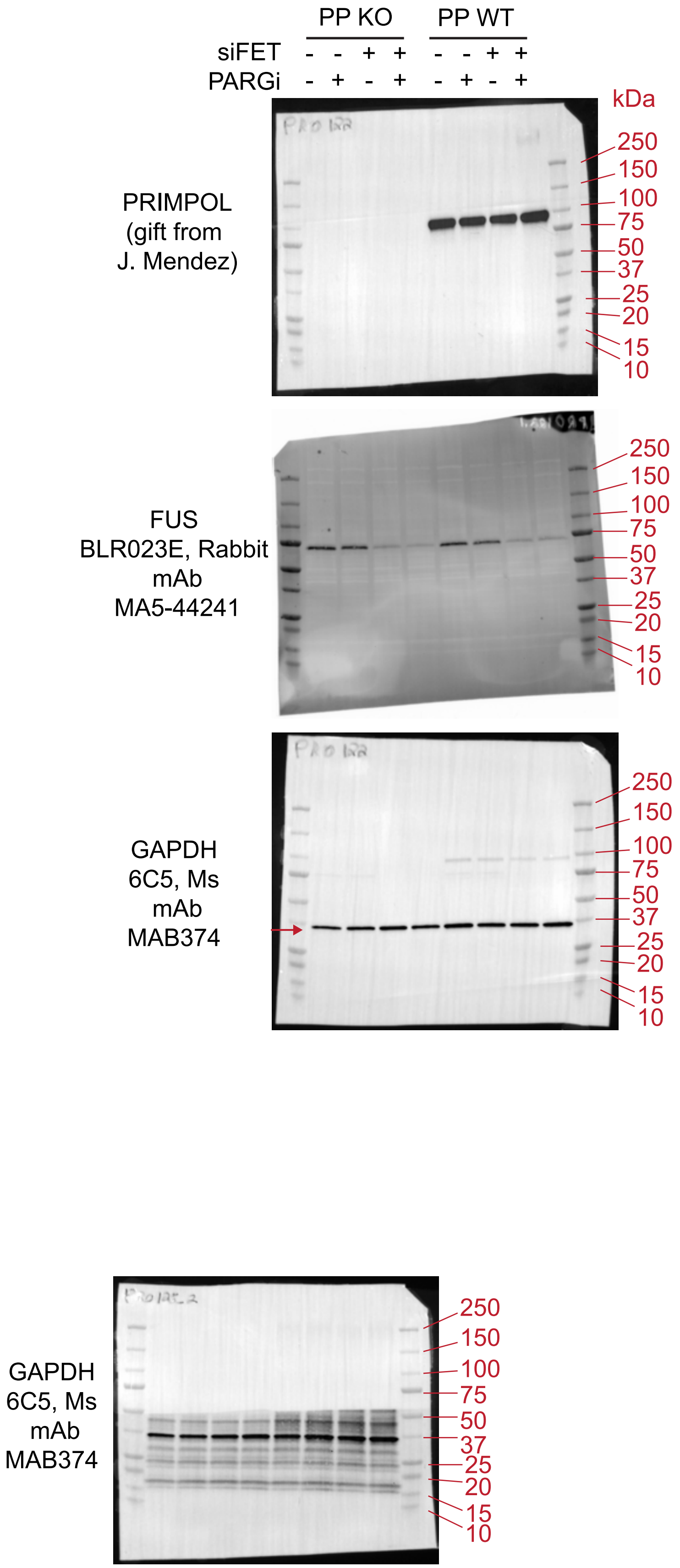

Supplementary Figure 5d

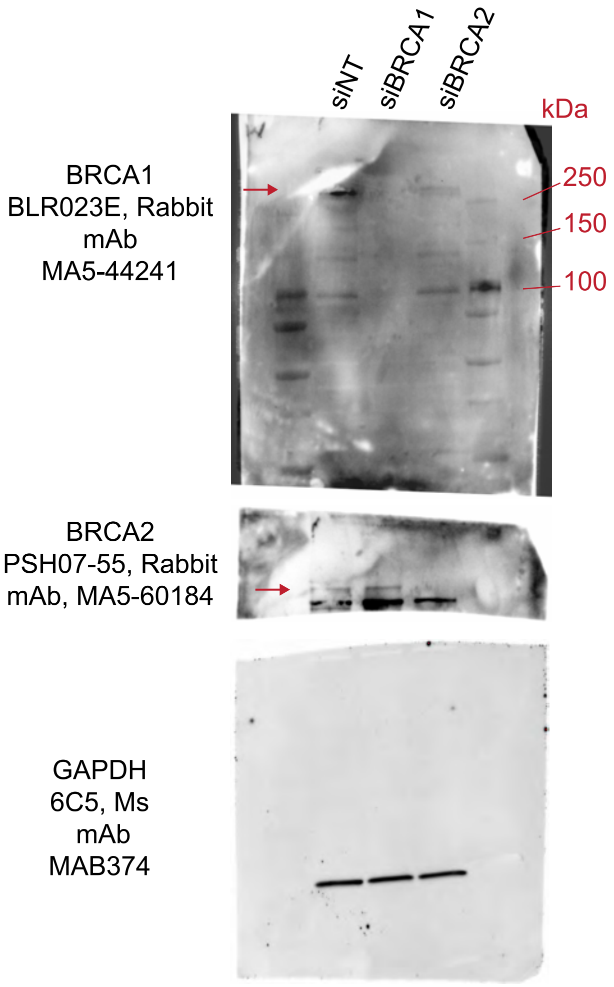


Supplementary Figure 5i

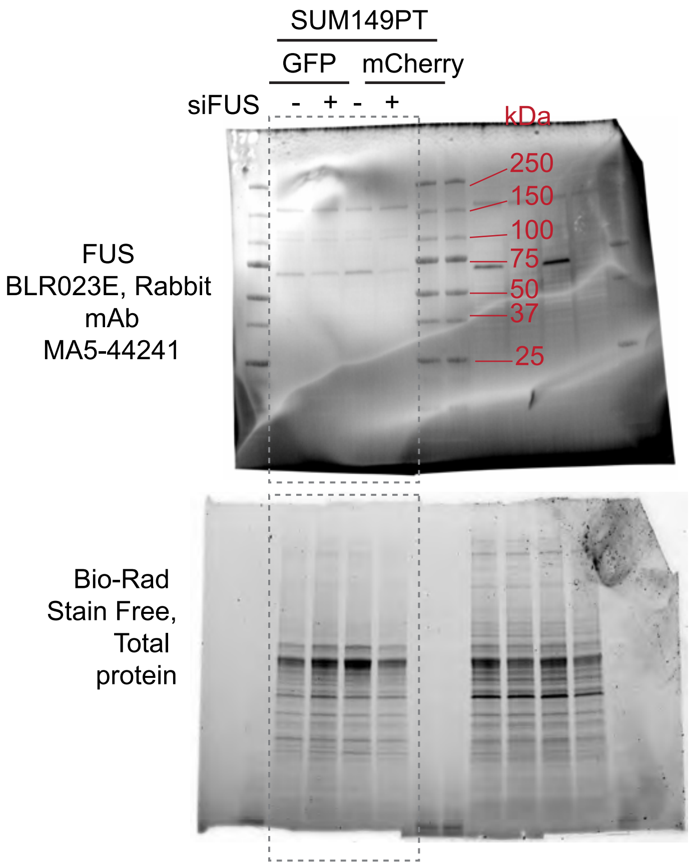


Supplementary Figure 5h

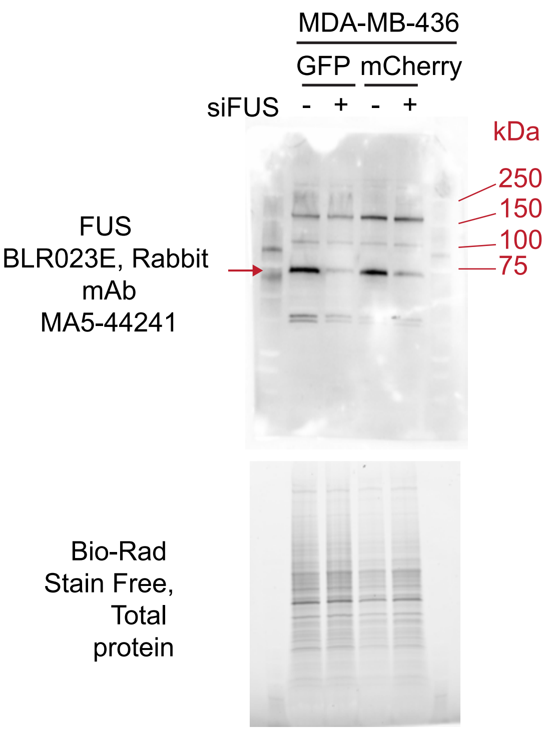

Supplementary Figure 2a

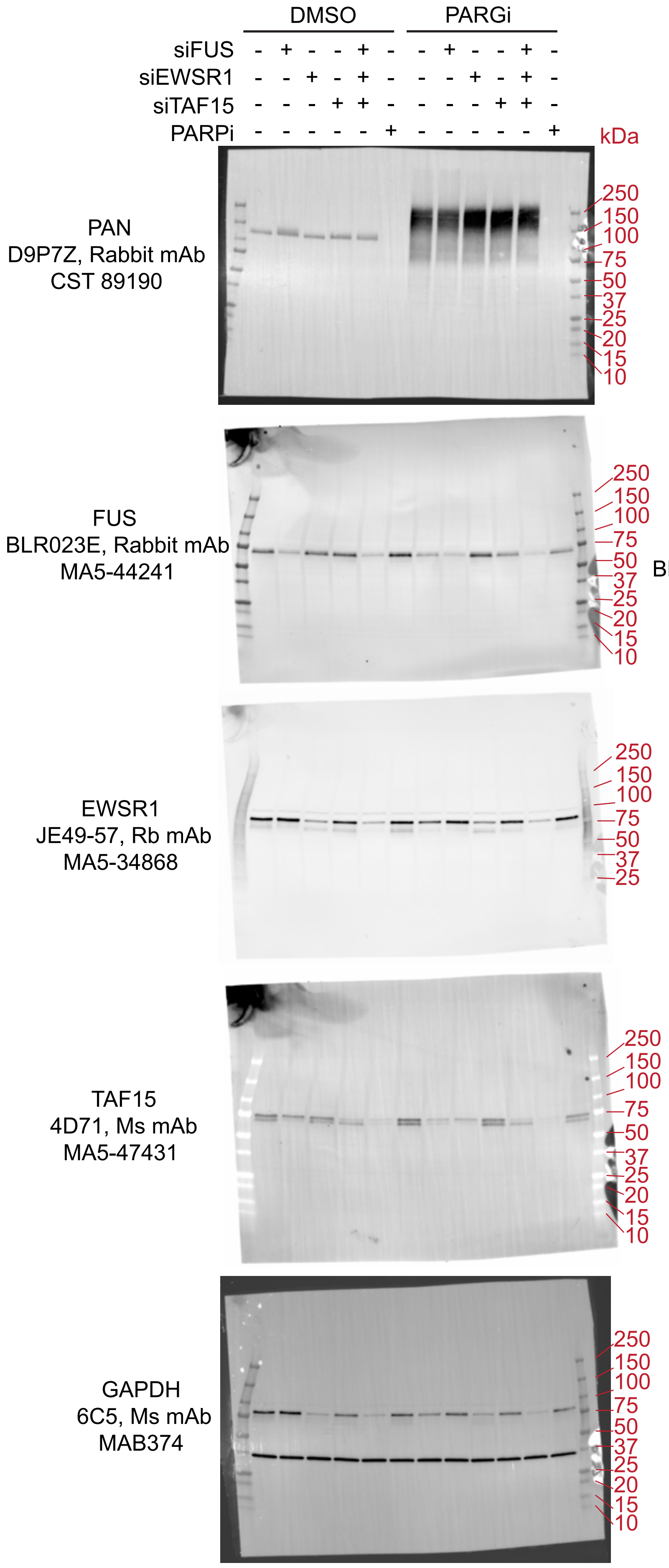


Supplementary Figure 2b

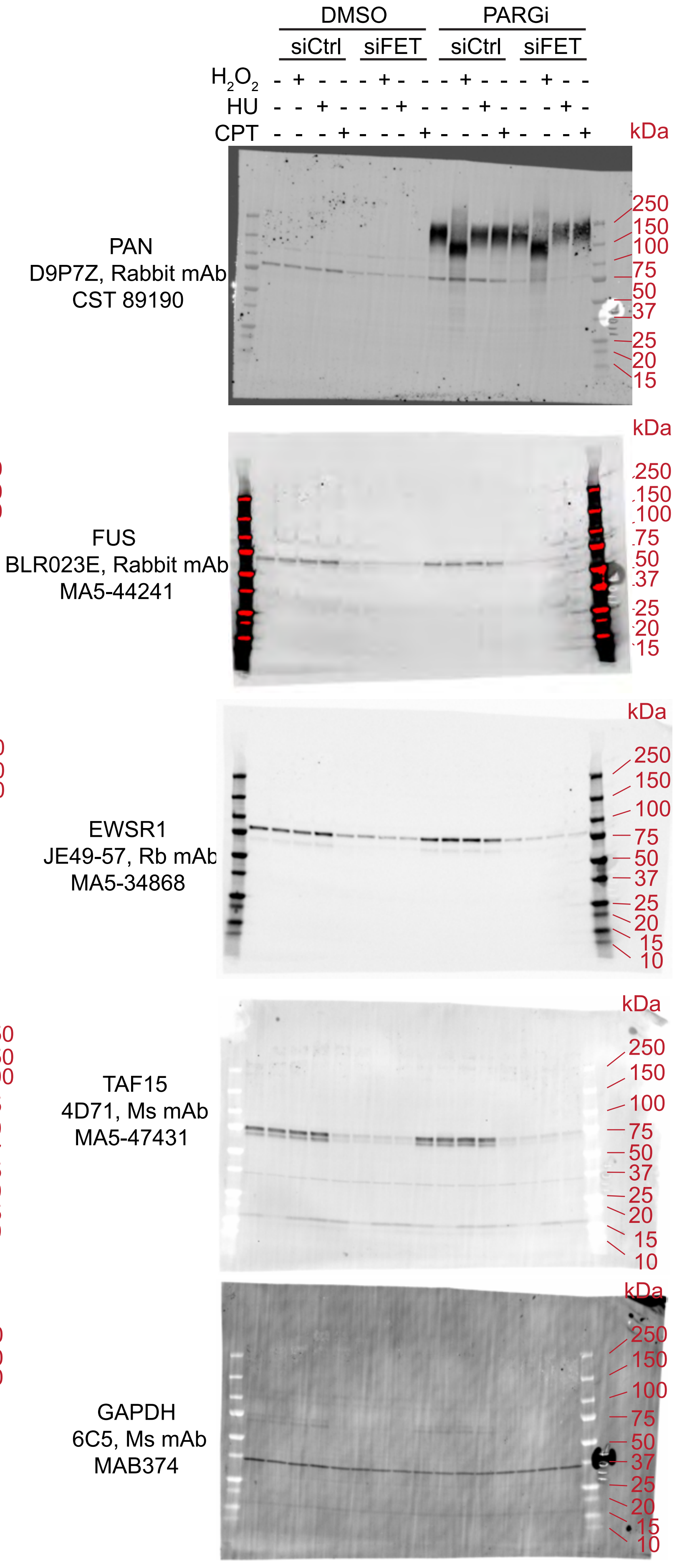
